# Learning millisecond protein dynamics from what is missing in NMR spectra

**DOI:** 10.1101/2025.03.19.642801

**Authors:** Hannah K. Wayment-Steele, Gina El Nesr, Ramith Hettiarachchi, Adedolapo M. Ojoawo, Hasindu Kariyawasam, Sergey Ovchinnikov, Dorothee Kern

## Abstract

Many proteins’ biological functions rely on interconversions between multiple conformations occurring at micro-to millisecond (µs-ms) timescales. A lack of standardized, large-scale experimental data has hindered obtaining a more predictive understanding of these motions. After curating >100 Nuclear Magnetic Resonance (NMR) relaxation datasets, we realized an observable for µs-ms dynamics might be hiding in plain sight. Millisecond dynamics can cause NMR signals to broaden beyond detection, leaving some residues not assigned in the chemical shift datasets of ~10,000 proteins deposited in the Biological Magnetic Resonance Data Bank (BMRB)^1^. We made the bold assumption that residues missing assignments are exchange-broadened due to µs-ms motions and trained various deep learning models to predict missing assignments. Strikingly, these models also predict exchange measured via NMR relaxation experiments, indicative of µs-ms dynamics. The best of these models, which we named Dyna-1, leverages an intermediate layer of the multimodal language model ESM-3^2^. Notably, dynamics directly linked to biological function, including enzyme catalysis and ligand binding, are particularly well predicted by Dyna-1, which parallels our findings that residues experiencing µs-ms exchange are more conserved. We anticipate the datasets and models presented here will be transformative in unlocking the common language of dynamics and function.

## Introduction

The functions of proteins often depend on their ability to interconvert between multiple conformations^3^. Our ability to understand fundamental biological mechanisms would be significantly improved if we had greater ability to predict multiple conformations and the timescales at which they interconvert. Computationally modelling the motions of proteins has been a longstanding goal since the first molecular dynamics (MD) simulations of proteins^4^. However, our predictive power for protein motions pales in comparison to our predictive power for single structures. AlphaFold-2 (AF2)^5^ demonstrated that the experimental data available in the Protein Data Bank (PDB)^6^, coupled with large-scale sequencing data and modern deep-learning architectures, could achieve unprecedented accuracy in structure prediction. However, the task of predicting *protein dynamics* lacks what the task of predicting single structures had: large-scale, standardized benchmarks of experimental observables. Any computational method, be it deep-learning-based or simulation-based, currently faces a paucity of standardized experimental data available on dynamics to evaluate on, let alone train from (**Fig. 1a**).

**Fig. 1.**
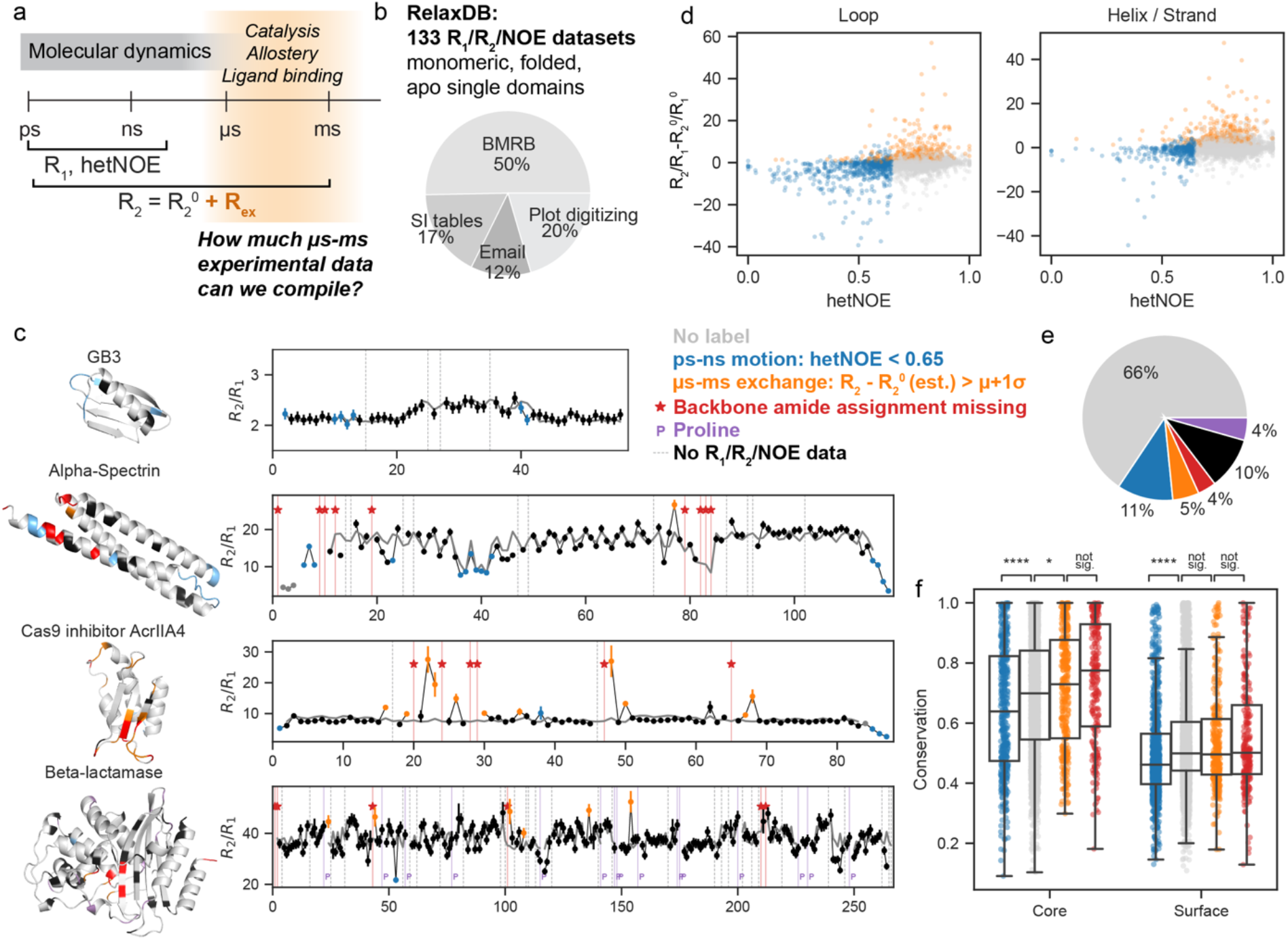
A benchmark of 133 protein NMR backbone relaxation measurements demonstrates residues with ps-ns motion are less conserved, and residues with µs-ms exchange are more conserved. (a) Many biological processes occur at a µs-ms time regime, yet experimental data in this regime have not existed at a sufficient scale for machine learning. NMR R_1_, R_2_ and heteronuclear NOE (hetNOE) measure ps to low ns motion, while millisecond timescale exchange causes increased R_2_ (R_ex_). (b) The “RelaxDB” benchmark we curated consists of 133 single-domain ^15^N protein backbone datasets with associated labels for the nature of motion present at each residue. Pie chart contains the origin of the data in RelaxDB. (c) Example raw R_2_/R_1_ data (black) and comparison to calculation from rigid tumbling using HYDRONMR (grey line)^48,49^ to obtain labels. Labels are shown in the different colors in legend and plotted onto the AF2 predicted structures. Error bars on R_2_/R_1_ data are calculated as the maximum of uncertainties reported by authors or uncertainties estimated from the data (see Methods). (d) Distribution of hetNOE vs. calculated change in R_2_/R_1_ between experiment and rigid tumbling calculation. Blue: ps-ns motion. Orange: µs-ms exchange. Grey: no label. (e) Distribution of RelaxDB labels. (f) Calculating sequence conservation for these residues demonstrates that residues with µs-ms exchange are more conserved than residues with no assigned label, which in turn are more conserved than residues with ps-ns motion. Residues with missing backbone amide assignments are no different in conservation than residues with µs-ms exchange. n=12,584 residues. Box plots depict median and 25/75% interquartile range, whiskers = 1.5 *interquartile range. Statistical comparisons by two-tailed independent-samples t-test; no multiple comparisons adjustment. *: 0.01 ≤ p ≤ 0.05; ****: p ≤ 0.0001.

We focus here on µs-ms dynamics, since it is well established that motions on this timescale are directly linked to biological function^7^ and are often concerted in nature^8^. Where can we find experimental data on µs-ms dynamics? The primary source of structural data used in deep learning, the structure information in the PDB, lacks time information. Time-resolved X-ray diffraction and cryo-electron microscopy methods are gaining increasing traction, but processing these datasets requires nontrivially deconvolving signal to identify multiple conformations. In contrast, the last 50 years of methods development in NMR has led to experiments in which the presence of µs-ms dynamics *creates* signal in the form the observable *R*_*ex*_. R_ex_ is an increase in R_2_ relaxation times that unambiguously indicates that the monitored atom is experiencing changes in its chemical environment at the µs-ms timescale^9,10^.

We wished to address the lack of experimental dynamics benchmarks by curating standardized datasets of R_ex_ measurements on backbone amides. In analyzing the resulting “RelaxDB” dataset of 133 proteins, we found that residues with µs-ms exchange were most conserved. This suggested that with more experimental data on µs-ms dynamics, it might be feasible to use an evolutionary-guided deep learning approach, such as a protein language model, to predict µs-ms dynamics. This task—namely, can a model predict the presence of a concerted, millisecond-timescale process?—is distinct from other existing tasks: predicting backbone flexibility or order parameters^11,12^ or predicting multiple conformations without information on timescales^13,14^. External conditions such as temperature, pH and cofactors will change the rate and relative populations of such a process, but not the underlying process itself.

We were curious if the ~10,000 proteins with deposited NMR chemical shift information in the BMRB could provide a useful dynamics observable for training a model. We hypothesized that missing ^15^N backbone assignments are primarily caused by exchange-broadening due to µs-ms motions. We tested this key assumption by training deep learning models to predict which residues are missing assignments, and consequently evaluating if these models also have predictive power for measured R_ex_ for residues that are assigned. Our results demonstrate that signatures of millisecond-timescale dynamics are indeed learnable by modern deep learning approaches trained with these NMR data.

The developed model, Dyna-1, is able to predict biologically relevant µs-ms exchange as assessed on proteins characterized by many independent labs. Notably, our model led to reinterpretation of published NMR relaxation data in several ways. While investigating “false negatives” in the RelaxDB dataset, we noticed that some proteins interact with phosphate ions as part of their biological function, yet the NMR relaxation experiments were conducted in phosphate buffer leading to artificially increased R_2_ due to buffer binding. Dyna-1 did not predict exchange from such non-specific phosphate binding. Conversely, Dyna-1 did predict millisecond exchange that typical analysis of Carr-Purcell-Meiboom-Gill (CPMG) experiments missed, but more careful consideration of the existing NMR data validated. We anticipate Dyna-1 as well as the datasets used for training and evaluation will be transformative in better understanding how biomolecular dynamics leads to function.

## Results

### Residues with exchange are most conserved

To curate datasets of standardized R_ex_ measurements, we started with the experiment where R_ex_ is most directly measured: Carr-Purcell-Meiboom-Gill (CPMG) relaxation dispersion^9,10,15,16^. However, we could only find fewer than 10 sets of complete ^15^N backbone CPMG data either publicly available or from private correspondences. We next turned to the more abundant “R_1_/R_2_/hetNOE” (R_1_ relaxation, R_2_ relaxation, heteronuclear nuclear Overhauser enhancement) experiments, which provide information on backbone pico-nanosecond (ps-ns) motions and µs-ms exchange. We compiled data on 163 distinct protein domains (**Fig. 1b**), the earliest of which were published in 1990^17^. Because R_ex_ reports on any process causing a change in chemical environment, which can include multimerization or the binding-unbinding of a ligand, we only included datasets of apo proteins with no cofactors present, where the authors had verified a monomeric sample. We also excluded intrinsically disordered proteins. We devised a framework to robustly and systematically label residues with exchange while accounting for anisotropic effects (see Methods).

**Fig. 1c** depicts the degree to which our framework accounts for anisotropic effects for four example proteins from RelaxDB. The protein GB3 is small yet has pronounced anisotropy in its R_2_/R_1_ arising from its helix; it was a system first used to demonstrate that anisotropic tumbling leads to heightened R_2_/R_1_^18^. This “zigzag” pattern due to differing directions of backbone N-H amide bond vectors in a helix is also evident in the highly-anisotropic helical bundle of alpha spectrin^19^. Our fitting process discerns between anisotropic data features and increases in R_2_/R_1_ due to µs-ms exchange, exemplified in AcrIIA4 Cas9 inhibitor^20^ and beta-lactamase^21^. The original 163 proteins were reduced to 133 as some proteins had extensive missing data or ps-ns motion indicative of disordered regions and could not be reliably fit with this workflow (see Methods). All datasets are in **Extended Data Fig. 1**.

**Fig. 1d** depicts the distribution of all residues from these proteins along the two experimental observables defining the presence of different timescales of motion: namely, dR_2_/R_1_ (see Methods) for µs-ms exchange, and hetNOE for ps-ns motion. As expected, many residues with low hetNOE have a dR_2_/R_1_ less than zero, as motions at the ps-ns timescale lead to both a lower hetNOE and a reduced R_2_ value. A few residues have both fast motions detected via hetNOE and elevated dR_2_/R_1,_ indicative of µs-ms exchange. If anything, we anticipate that this method of looking at elevated R_2_/R_1_ is under-estimating µs-ms exchange for residues where motions are present at more than one timescale, as µs-ms exchange in elevating R_2_ and ps-ns motion decreasing R_2_ would cancel.

Despite comprising only 133 proteins, the RelaxDB benchmark represents an unprecedented amount of experimental information on dynamics in one place. **Fig. 1e** depicts the overall distribution of different labels assigned per residue. With this curated dataset in hand, we were interested in seeing if there were any eminent relationships between sequence and dynamics. When calculating sequence conservation, we noticed a striking trend: the residues with µs-ms exchange were more conserved than residues with no designated label, which in turn were more conserved than residues with ps-ns motion. Overall, solvent-exposed residues are less conserved than non-solvent-exposed residues (**Fig. 1f**). Residues involved in µs-ms processes are often involved in functions such as allosteric signaling, catalysis, or ligand binding. Nevertheless, we were surprised that this could be quantitatively detected in as few as 133 proteins. Further analysis revealed that the predicted local distance difference test (pLDDT) measure from AF2 is lowest for residues with ps-ns motion and shows little discrepancy between residues with no motion and µs-ms exchange (**Extended Data Fig. 2**).

Though we hoped RelaxDB would be useful for evaluating models, any model trained with only 133 proteins is unlikely to generalize. In looking at the data, we realized that our initial curation of the data lacked a key observable that could also give information on µs-ms dynamics. Some residues in the proteins of RelaxDB could not be assigned a label because they did not have measured R_1_/R_2_/hetNOE values. These residues lacking relaxation data fall into two fundamentally different categories. In the first category, a residue is assigned, but the authors did not report R_1_/R_2_/hetNOE values for a variety of reasons: possibly the peaks were too overlapped to fit exponentials reliably or an exponential did not fit well, for instance due to weak signal intensity. In the second category, a residue is missing R_1_/R_2_/hetNOE values because the residue was unable to be assigned in the first place. Crucially, residues can be missing from chemical shift assignments if they are “exchange-broadened”, i.e., they are experiencing exchange between two states where the chemical shift difference of the two states is comparable to the frequency at which the residue is interconverting. Therefore, we identified the corresponding published assignments for each protein and created labels to distinguish whether residues without R_1_/R_2_/hetNOE data were assigned or not. We hypothesized that if the residues missing assignments were also reporting on residues with µs-ms exchange, they would show similar trends in conservation to residues with R_ex_. Indeed, sequence conservation is statistically indistinguishable between residues with R_ex_ and residues with missing assignments (**Fig. 1f**). Fundamentally, whether a residue is not assigned because it is exchange-broadened or can be assigned but displays R_ex_ is dependent on the chemical shift difference of its multiple states relative to their interconversion rate.

### Curating proxy labels for exchange

We realized that these missing assignments were present at orders-of-magnitude higher counts than in just the relaxation measurements (**Fig. 2a**); the Biological Magnetic Resonance Bank (BMRB) contains ~12,000 proteins with deposited chemical shifts. Could a model trained on all the proteins in the BMRB, where the label is “is this residue assigned or not?” generalize to predicting µs-ms exchange? The key assumption such a model would be leveraging is that backbone amide assignments are primarily missing due to exchange, as opposed to incomplete assignments or other reasons. Of course, residues that are completely exchange-broadened represent a subset of all residues with exchange. If we use residue counts in RelaxDB as a rough proxy for the relative abundances of detectable R_ex_ versus missing assignments, we observe that 5% of residues have elevated R_2_/R_1_, whereas 4% have missing assignments (**Fig. 1e**). This rough estimate concludes that nearly half of the residues with exchange have missing assignments, which certainly is not all, but also not negligible.

**Fig. 2.**
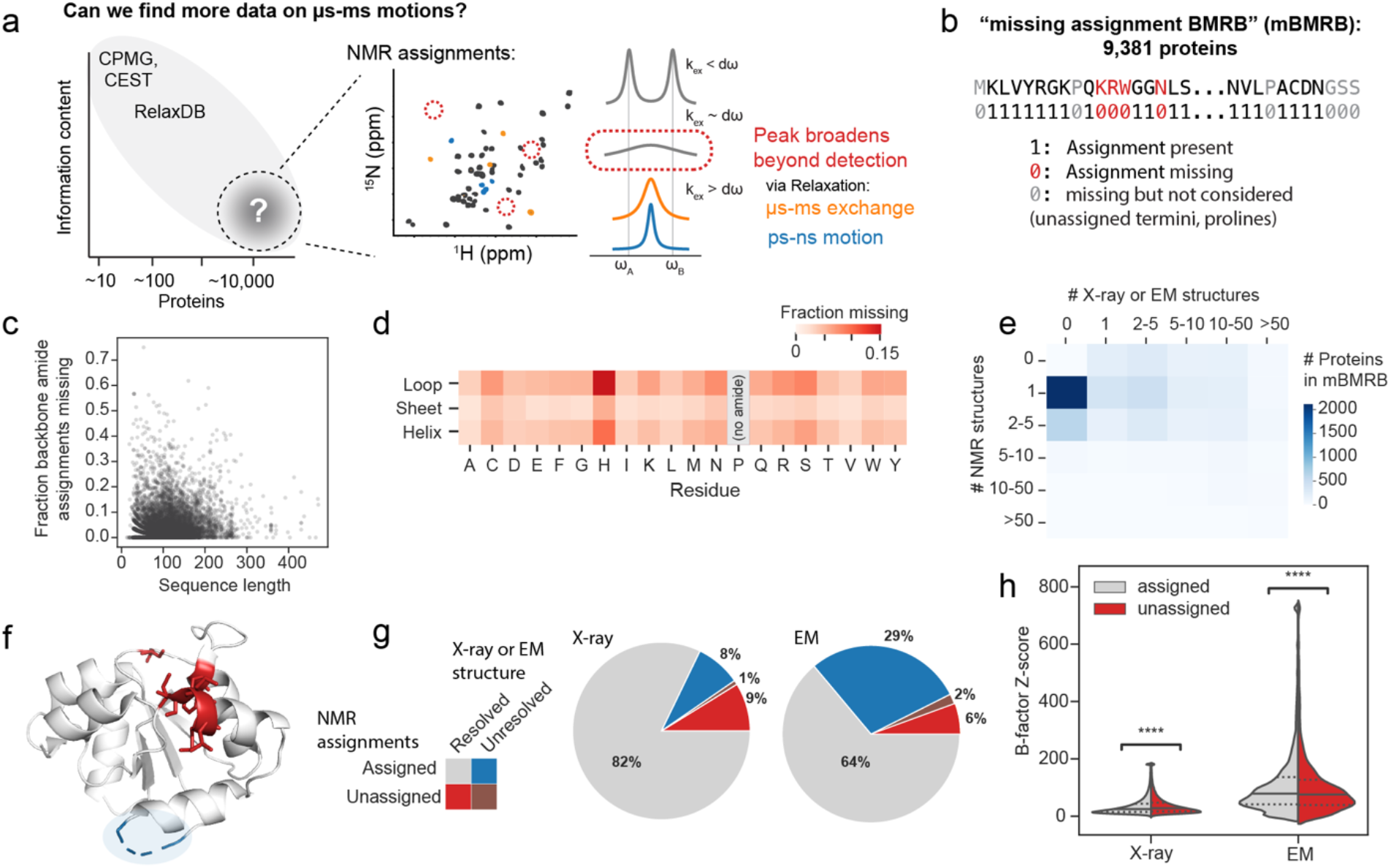
Missing NMR amide backbone assignments in BMRB used as label for µs-ms exchange. (a) In seeking more experimental data for µs-ms dynamics, we realized that data on roughly 10,000 proteins exists in the form of missing backbone amide assignments in the BMRB. Theoretical NMR line-shapes for 2-site exchange on different timescales shown. (b) Scheme for curating the “missing assignment BMRB” (mBMRB) dataset. (c) Fraction backbone amides missing vs. sequence length, not considering prolines or unassigned termini. (d) Fraction of backbone amides missing, disaggregated by amino acid and secondary structure type, in structures predicted by ESMFold^22^. (e) The majority of proteins in the BMRB have only one NMR structure and no X-ray or EM structure. (f) Residues with missing NMR peaks are distinct from residues missing in X-ray and EM structures, as exemplified by the MAP kinase binding domain of DUSP 16 (BMRB:19330, PDB:3TG3). Residues with missing assignments are in red, residues with missing X-ray density are in blue. (g) Fraction of residues assigned/unassigned in the mBMRB compared with resolved/unresolved in X-ray or EM structures shows little overlap. (h) B-factors of residues with present or missing NMR peaks have similar distributions. Solid lines: median, dotted lines: 25/75% percentile. In 2(g,h): n=71,704 protein datasets, n=306,842 residues. Statistical comparisons by two-tailed independent-samples t-test; no multiple comparisons adjustment. ****: p ≤ 0.0001.

We curated a new dataset: the mBMRB. In this dataset, we removed proteins that might have systematic, non-exchange reasons for assignments being missing (see Methods, **Extended Data Fig. 3**). This resulted in a dataset containing 9,381 proteins, almost two orders of magnitude more than in RelaxDB (**Fig. 2b**). This dataset comprises “positives” (for µs-ms dynamics), which are missing assignments curated as carefully as possible, but of course the “negatives” (no µs-ms dynamics) in this dataset are not unambiguously so: many residues are assignable while also experiencing exchange. Nevertheless, after applying these filters, we observed that a substantial fraction of proteins across the BMRB had significant numbers of non-proline and non-termini missing assignments (**Fig. 2c**). Disaggregating residues by identity and secondary structure type (calculated from structures predicted by ESMFold^22^) reveals that missing assignments span across both loops and structured elements for all amino acid types, though more frequently in loops (**Fig. 2d**). We note that while there will doubtlessly be noise in these labels arising from mis-assignment or other errors, there is well-established literature showing that in certain regimes of machine learning, noise can actually assist model convergence as a form of regularization^23,24^.

We next compared the information in these “missing assignment” labels to sources of heterogeneity and uncertainty present in other forms of publicly available structure data to inform on what models trained on the PDB such as AF2 may or may not have already been exposed to (see Methods). 45% of these proteins (2680/5906) had no X-ray or EM structures but had one or more NMR structures (**Fig. 2e**). For proteins that were characterized by X-ray and/or EM and assigned by NMR, residues with missing assignments appear distinct from residues unresolved in X-ray or EM structure models (**Fig. 2f,g**). Of the residues that are resolved, the distributions of B-factors are similar for both assigned and unassigned residues (**Fig. 2h**). In summary, our mBMRB dataset contains unique information on thousands of proteins in comparison to information present in X-ray and EM structures in the PDB.

### Predicting missing NMR assignments

We next tested if current deep learning models when trained with these curated NMR data have the capacity to predict which assignments are missing, and second if such learning could transfer to predictive power for µs-ms exchange. We created a train/validation/test data split, initially holding out any sequences with >80% sequence identity to the RelaxDB or RelaxDB-CPMG datasets to use later as higher quality experimental evaluations with labels of exchange (**Fig. 3a**). To contextualize model performance on this new task, we began by calculating a set of naïve baselines. Of these, the best performing model on the validation set used sequence, secondary structure, and SASA (see Methods, **Extended Data Fig. 4a**).

**Fig. 3.**
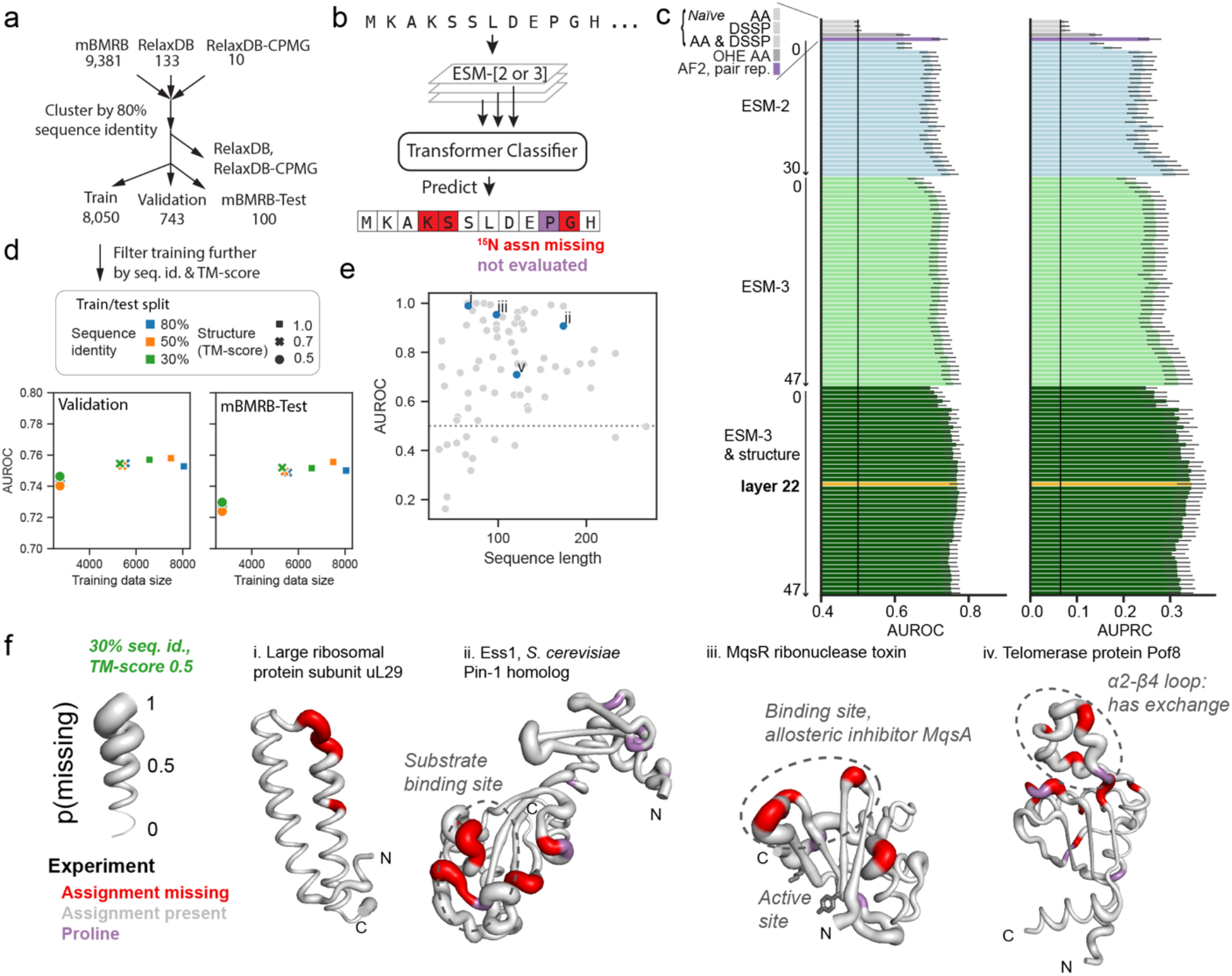
Deep learning has predictive power for the “missing assignment BMRB” (mBMRB). (a) Train/validation/test split scheme to train from the mBMRB while holding out 100 proteins from mBMRB as well as all RelaxDB and RelaxDB-CPMG data as test sets. (b) Schematic of deep learning architectures tested. (c) AUROC and AUPRC on validation set testing layers from ESM-2^22^ and ESM-3^2^ models, AlphaFold2^5^ pair representation, and baselines. AA: amino acid. DSSP: secondary structure assignment. OHE AA: one-hot encoded amino acid. Layer 22 was chosen as the best-performing model (Dyna-1). Error bars represent 95% confidence interval for AUROC and AUPRC evaluated over proteins in validation set. (d) Performance is not significantly impacted by removing sequence or structure homologues more stringently from training. (e) AUROC from Dyna-1, trained using the 30% sequence identity and 0.5 TM-score dataset split, on the mBMRB-Test set. Proteins in blue are shown in (f). (f) Examples of Dyna-1 predictions. Structures are generated with ESMFold from mBMRB sequence (BMRB entries 5977, 50787, 51503, and 50002 for i-iv, respectively), the thickness of the tube is the probability missing as predicted by Dyna-1. Residues are colored for experimental data as follows: red: missing backbone assignment, grey: assignment present, purple: proline (no data).

We were curious to explore the predictive power of existing deep learning models on this task, since AF2^5^ and ESM-2^22^ have shown predictive capabilities beyond the tasks for which they were trained. For example, the protein language model ESM-2 was trained in the self-supervised task of masked language modeling yet has predictive power for secondary structure, 3D structure, and functional labels. In this “transfer learning” paradigm, extracted representations from pretrained models are used to train new models with different datasets on related downstream tasks^25^. We designed architectures to test structure-focused representations from AF2^5^ pair representations (“AF2, pair rep.”), sequence-only representations in ESM-2^22^, and interactions between sequence and structure by using the multimodal language model ESM-3^2^ (see Methods, **Fig. 3b, Extended Data Fig. 5**). We first compared these architectures by training with our curated NMR dataset (mBMRB) at 80% sequence identity cutoff; after assessing the models’ predictive capability on this novel task given a majority of the available data, we then retrained models with a more stringently filtered training set (see below).

We observed that all three of these pretrained models out-performed other baseline models (**Fig. 3c**). The best-performing model was layer 22 of ESM-3 if provided both sequence and structure models as input, with an AUROC of 0.77. Intriguingly, the last layer of ESM-2 model achieved an AUROC of 0.75, and the AF2-pair model, which did not use MSAs, and rather only structure input, achieved 0.71. This suggests that both sequence and structure information provide utility in this new task. Performance from all ESM-2 and ESM-3 layers tested are depicted in **Fig. 3c**. AUPRC (**Fig. 3c**, right) mirrors AUROC values. We selected based on AUPRC since it is slightly more discriminative than AUROC. AUPRC visualizations for all subsequent per-protein comparisons discussed are included in **Extended Data Fig. 6**.

We tested the effect of sequence and structure similarity in the training data on performance by training with more stringent training splits: we tested 50% and 30% sequence identity cutoffs and structure-based similarity cutoffs of TM-score at 0.5 and 0.7. These significantly reduced the size of the training set, with the smallest including only ~2700 proteins. The most stringent training split decreased the AUROC of Dyna-1 on the validation set to 0.74 (**Fig. 3d**), and comparably on the remaining evaluation datasets in this work (**Extended Data Fig. 6**). We did not observe any individual proteins with significant jumps in AUROC between our most stringent training set (30% sequence identity, 0.5 TM-score cutoff) and our most lenient (80% sequence identity, 1.0 TM-score cutoff), which would have indicated model memorization (**Extended Data Fig. 5d**). We note that model performance does not improve significantly when given almost three-fold more data (cf. **Extended Data Fig. 6**). The final model leverages the embeddings of layer 22 of ESM-3, uses both sequence and structure as inputs, and was trained with the most stringent data split (30% sequence identity, 0.5 TM-score cutoff).

**Fig. 3e** depicts AUROC per protein for those with one or more missing assignments in the mBMRB-Test dataset. **Fig. 3f** depicts the structures of some of the best-performing proteins in the test set with tube sized by the output of Dyna-1, i.e., the predicted probability each residue is missing, “p(missing)”, in comparison to residues with missing assignments (red). The missing assignments in example (iii) are related to the function of this protein; in ribonuclease toxin MqsR, loop β2-β3 is known to have exchange, and its flexibility is reported to be related to its ability to bind mRNA^26^. The antitoxin MqsA acts on this protein not by directly inhibiting its active site, but rather by sequestering the β2-β3 loop and prohibiting its motion, which allows the ribonuclease to show broad non-specificity towards a variety of mRNA substrates^26^.

Though these results appeared promising, missing assignments are an incomplete indicator of µs-ms dynamics, since residues can experience exchange broadening (R_ex_) that does not necessarily preclude their assignment. Hence, “false positives” in this data cannot be held as ground truth. This is exemplified in (iv), the telomerase protein Pof8^27^. Dyna-1 predicts high p(missing) for the α2-β4 loop, yet most of these residues are assigned. However, the authors described that assigning the α2-β4 loop and analyzing subsequent relaxation data for the loop was difficult “due to severe line broadening”^27^, a clear indicator of µs-ms exchange. We therefore needed to evaluate Dyna-1’s predictions using the more-informative relaxation data in RelaxDB to gain a better sense of the model’s predictive capability for µs-ms dynamics. This next evaluation also tests if model learning can “transfer” beyond only predicting residues that are exchange-broadened to the extent that precludes their assignment to residues that are assignable, but have exchange detectable via relaxation.

### Dyna-1 predicts µs-ms exchange

We found that when evaluating only missing assignments (**Fig. 4ai**), representative models achieved comparable performance to the mBMRB validation and test sets. To test if the model indeed generalized beyond the data with which it was trained, we held out missing assignments and asked how well it could predict which assigned residues have exchange (**Fig. 4aii**). Performance does decrease across all models compared to the task of predicting missing assignments, but AF2-pair, ESM-2, and ESM-3-based models achieved AUROC well above the controls, indicating that the models can generalize learning from predicting missing assignments (e.g. p(missing)) to predicting exchange within assigned residues (e.g. p(exchange)). To evaluate the performance of the model to predict µs-ms exchange as manifested in both forms, we consider both missing assignments and residues with measured µs-ms exchange in RelaxDB (**Fig. 4aiii**). Out of the representative models, layer 22 of ESM-3 given both sequence and structure again performed the best in this evaluation, same as in the mBMRB validation set. We named this final model “Dyna-1”.

**Fig. 4.**
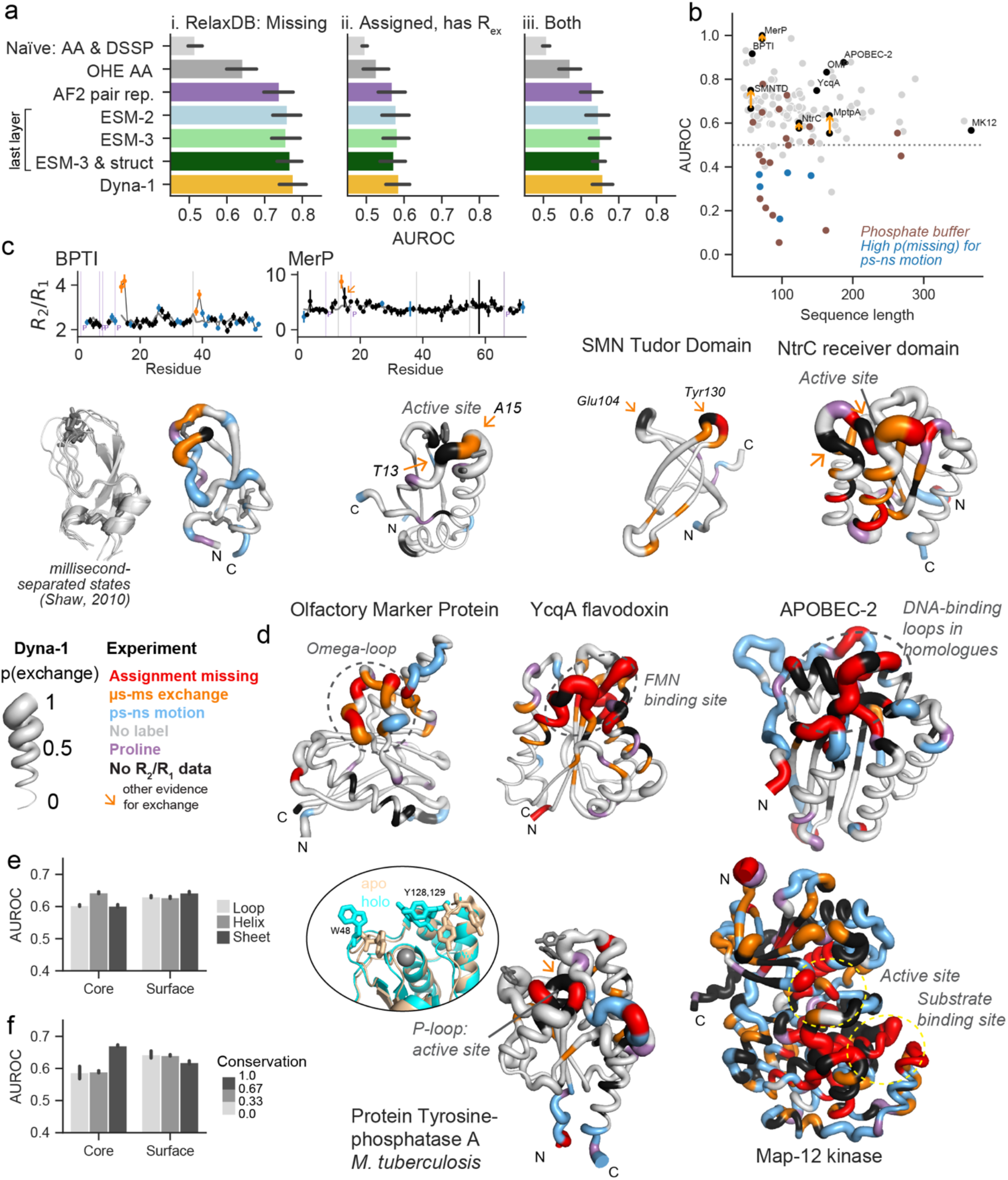
Dyna-1 predicts biologically-relevant micro-millisecond dynamics in NMR relaxation data. (a) Representative models evaluated on (i) missing assignments in RelaxDB, (ii) residues with assigned R_ex_, and (iii) both missing and R_ex_. Error bars represent 95% confidence interval for mean evaluated over proteins in RelaxDB. Error bars represent 95% confidence interval for mean AUROC evaluated over proteins in RelaxDB (n=112). (b) AUROC per protein vs. sequence length for Dyna-1 in RelaxDB for both missing assignments and residues with R_ex_. Only shown are proteins with 2 or more residues with missing assignment or exchange. Brown: proteins with exchange from phosphate buffer. Blue: proteins with high p(exchange) primarily in regions with ps-ns motions (cf. Extended Data Fig. 7). Orange arrows: proteins where follow-up on residues with no relaxation data resulted in increased AUROC (see c,d). (c) Dyna-1 predictions on smaller proteins offer an opportunity for detailed comparisons to experimental data. BPTI agrees with experimental R_2_/R_1_ data^31^ (top) and heterogeneity among kinetically-distinct substates in molecular dynamics^32^ (left). Mercuric Transport Protein, SMN Tudor Domain, and NtrC receiver domain offer prospective tests of residues with high p(exchange) for which no data was available (black), but other evidence for R_ex_ (orange arrow). R_2_/R_1_ error bars are from the original curated datasets. (d) Representative Dyna-1 predictions for larger proteins. Biologically-relevant features are annotated in grey. The Dyna-1 prediction for MptpA, a phosphatase from *M. tuberculosis*, recapitulates measurements from NMR as well as structural heterogeneity in sidechains regulating specificity (W48, Y128, Y129) (apo: 2LUO, holo: 1U2P). (e) AUROC for residues binned by core and surface designation and secondary structure type indicates that Dyna-1 has the most predictive power in core helices and surface loops. (f) AUROC binned by core/surface location and sequence conservation indicate that Dyna-1 has the most predictive power for more conserved residues. In (e,f), bars represent mean; error bars represent standard error evaluated via bootstrapping (n = 13,810 residues).

**Fig. 4b** depicts AUROC for all proteins in RelaxDB vs. sequence length. We first noticed that there were numerous proteins in RelaxDB with an AUROC of <0.5, i.e. worse than a random predictor. By carefully reading the corresponding NMR papers, we learned that some of these were DNA- or RNA-binding proteins measured in phosphate buffer, and that the unpredicted but experimentally reported exchange often occurred at positively-charged surface residues (**Extended Data Fig. 7**). Several publications directly state this phenomena as an artefact about either their own or others’ datasets being a limitation to interpreting measured exchange in the system they investigate.^28,29^ We annotated these low-performing datasets further and found that out of the 22 proteins with AUROC ≤ 0.5, 10 of these (45%) are proteins that during their biological function would be expected to interact with phosphate groups (i.e. DNA- or RNA-binding proteins or signaling domains), yet were experimentally characterized in phosphate buffer. Removing these proteins from RelaxDB increased Dyna-1’s overall performance from an AUROC of 0.63 to 0.66.

**Figs. 4c,e** depict representative proteins where structures are sized by Dyna-1 predicted p(exchange) and colored by experimental data for comparison. In Bovine Pancreatic Trypsin Inhibitor (BPTI), Cys14, Lys15, Cys38, and Arg39 are known to undergo millisecond transitions from disulfide bond isomerization.^30,31^ Promisingly for our automated labeling scheme, these four residues are also labeled as having µs-ms exchange from relaxation data in ref. ^31^ (depicted in **Fig. 4c**, top). Shaw et al. identified 5 kinetically distinct states on the micro-millisecond timescales via molecular dynamics simulations^32^ (depicted in **Fig. 4c**, left). Strikingly, Dyna-1 predicts high p(exchange) for the two loops containing these residues (**Fig. 4c**, right), to achieve an AUROC of 0.92 for predicting millisecond exchange in BPTI. While investigating Dyna-1 predictions in RelaxDB and revisiting original datasets, we found several residues with no relaxation data yet other evidence exists for exchange, underscoring Dyna-1’s predictive power (see supporting information, orange arrows in **Fig. 4c,d**).

In assessing what motifs Dyna-1 was able to predict in larger proteins (**Fig. 4d**), we were intrigued to see that many of the dynamic motifs it correctly predicts also have biological relevance. In flavodoxin YcqA, Dyna-1 predicts high exchange in the loop and helical regions that correspond to the FMN ligand-binding site. In Olfactory Marker Protein from humans, Dyna-1 predicts high p(exchange) in the omega-loop, which is described in ref. ^33^ to have substantial R_ex_ and is highly conserved. More recent studies indicate that the omega loop serves as a nuclear export signal^34^. In APOBEC-2, Dyna-1 predicts high p(exchange) in a region that in structure homologues is known to control the specificity of DNA substrates^35^. For Tyrosine-phosphatase A from *M. tuberculosis* (MptpA)^36^, Dyna-1 has high p(exchange) in loops containing residues with differing orientations between the apo and holo structures (apo: 2LUO, holo: 1U2P), and which are postulated to be involved in regulating enzyme specificity^36^. To test our hypothesis that Dyna-1 was performing better in predicting conserved motions, we calculated AUROC across different sets of residues categorized by secondary structure, conservation score, and core and surface designation. Indeed, we found that Dyna-1 had the most predictive power for more conserved residues in the core, supporting our observations on individual proteins (**Fig. 4e,f**).

Dyna-1 predictions in the MAP kinase p38 gamma (MK12) are indicative of room for future improvement. For MK12, we used experimental labels derived from the authors’ model-free fitting to multiple field strengths^37^. MK12 is a highly dynamic protein with ps-ns and µs-ms processes occurring throughout its structure, and p(exchange) predicted by Dyna-1 was high across the whole protein. Improving discrimination between ps-ns motions and µs-ms exchange is the subject of future work. We identified numerous proteins where Dyna-1’s highest p(exchange) is in regions with ps-ns motion (blue in **Fig. 4b**, structures in **Extended Data Fig. 7c**). One challenge in interpreting these is that residues can experience both ps-ns motions and µs-ms exchange, and single field strength measurements for R1 and R2 are insufficient to experimentally quantify motions at two timescales simultaneously. We expect that if dynamics is present at more than one timescale, because µs-ms exchange elevates R_2_ and ps-ns motion decreases R_2_, our current experimental labels for those likely underestimate R_ex_.

### Dyna-1 reveals overlooked exchange

Due to these challenges in discerning different types of dynamics in single-field-strength R_1_/R_2_/hetNOE experiments, we lastly tested Dyna-1 on proteins with µs-ms exchange directly characterized by ^15^N CPMG relaxation dispersion (RelaxDB-CPMG dataset, Fig. 5). CPMG experiments allow for directly measuring contribution from conformational exchange to R_2,eff_^9,15,16^. In brief, data are collected with increasingly short gaps between spin-echo pulses, which increasingly suppress contributions from exchange between states with different chemical shifts (R_ex_) to R_2_^10,15,16^. In typical data processing (orange trace, **Fig. 5a**), R_2,eff_ plateaus at a value that can be understood to be R_2_ in the absence of chemical exchange (R_2,inf_). The difference between R_2,eff_ at the lowest B_1_ field strength and R_2,inf_ is the R_ex_ contribution to R_2_ that was suppressed. We note that because our RelaxDB-CPMG dataset is curated with ^15^N relaxation data, our evaluations on the experimental data do not consider cases where R_ex_ is only seen for ^1^H caused by chemical shift differences only in ^1^H and not ^15^N, therefore underestimating other possible chemical exchange. Hence, Dyna-1 performance might be better than reported.

**Fig. 5.**
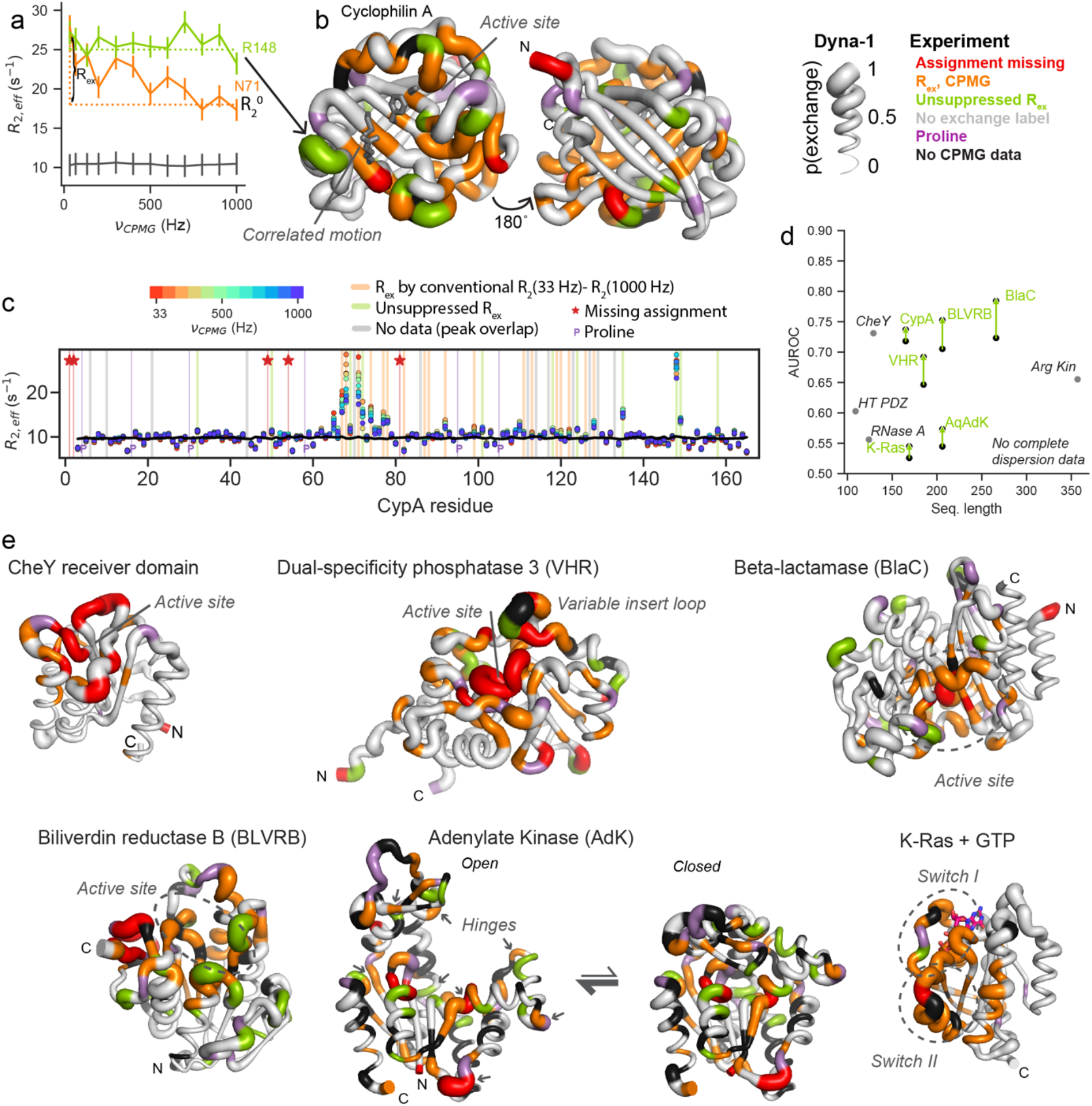
Dyna-1 even predicts exchange in residues that conventional CPMG data analysis misses. (a) Example ^15^N CPMG data for 3 residues from Cyclophilin A (CypA)^40^. Error bars represent background noise in spectrum. R_ex_ is typically estimated via difference in R_2,eff_ at lowest field strength and R_2_^0^ value to which R_2,eff_ plateaus. However, plotting R_2,eff_ for Arg148 (green) demonstrates different behavior: R_2,eff_ remains high over all B_1_ field strengths used, indicative of exchange on the microsecond regime that is too fast to be suppressed with the CPMG field strengths used. (b) AF2 structure of CypA with tube thickness by Dyna-1 p(exchange). Legend coloring according to the experimental data is shown to the right. Dyna-1 predicts high exchange for Arg148. (c) We devised a data-processing method using R_2_ predicted by HYDRONMR to systematically assign residues with R_ex_ for residues where R_2,eff_ get suppressed by CPMG pulse train (conventional R_ex_, in orange) or residues with elevated, unsuppressed R_ex_ (green). Spectra depicted with representative data for CypA was collected at 25°C. (d) Incorporating labels for unsuppressed R_ex_ increases reported AUROC for 5/6 proteins where complete dispersion data was available for reprocessing (shown by green arrows). Also shown is the AUROC for proteins where we could not find dispersion data for all residues, and instead curated residue labels based on descriptions in literature. (e) Representative experimental data and Dyna-1 predictions for a selection of proteins with dynamics characterized with ^15^N CPMG. For Adenylate Kinase, structure predictions of the open and closed states from AF2 were used as input into Dyna-1.

We first evaluated Dyna-1 by creating labels for residues with exchange following this data processing. **Fig. 5b** depicts Cyclophilin A (CypA) sized by Dyna-1 prediction probabilities, where residues with statistically significant R_ex_ from ref. ^38^ are colored orange. Importantly, Dyna-1 predicted higher p(exchange) in the beta-sheet with a known concerted millisecond dynamics involving the sidechains of residues Ser99, Phe113, Met61, and Arg55^39^ (shown in grey in **Fig. 5b**). Note that the other beta-sheet on the back of CypA, is correctly predicted to have no exchange. We were curious about residue Arg148: Dyna-1 predicted high exchange probability, but Arg148 did not register as having significant R_ex_ as evaluated by the above method. When looking at the raw data, we found that Arg148 had high R_2,inf_ compared to other residues. We propose that Arg148 has exchange in a process that is too fast to be suppressed by the B_1_ fields used in the CPMG experiment for the given temperature. However, to confirm this requires a reasonable estimate for what R_2,inf_ should be if R_2,eff_ is never sufficiently suppressed by the CPMG field strength. We devised a new framework to estimate anisotropic effects from rigid tumbling to then identify unsuppressed R_ex_ (see Methods). **Fig. 5c** depicts CPMG data for CypA collected at 25°C and 600 MHz from a B_1_ field strength of 33 (red) to 1000 Hz (blue). Several residues have unsuppressed R_ex_ (in green) which conventional data processing would not have identified as having exchange.

We applied our new processing to 6 datasets where we had ^15^N CPMG dispersion data for all non-overlapped residues, including CypA^40^, beta-lactamase^21^, Adenylate Kinase from *Aquifex Aeolicus*^*41*^, Biliverdin reductase B^42^, K-Ras^43^, and Dual-specificity phosphatase 3^44^ (**Extended Data Fig. 8**). For all datasets, Dyna-1’s performance improved when considering residues with unsuppressed R_ex_ (**Fig. 5d**, structures in **Fig. 5e**). Dyna-1 also predicts unsuppressed R_ex_ in Biliverdin reductase B, a cellular redox regulator, where millisecond motions are implicated in efficient coenzyme engagement and catalysis^42^, and predicts concerted millisecond dynamics in the switch I and II regions of K-Ras that is well known to be crucial for its signaling activity^43^. We additionally analyzed 4 proteins where CPMG was performed but raw data was unavailable, and found that Dyna-1 had predictive power for residues other labs reported had exchange (**Extended Data Fig. 9**).

### Prospective experimental test of Dyna-1

To further validate the predictive capabilities of Dyna-1, we conducted a prospective experimental test after public release of the model and initial review of the manuscript. We selected proteins from the “mBMRB test” set to measure their dynamics, as they are already assigned but many do not have published relaxation data. Plotting the mean p(exchange) from each protein in the mBMRB test set vs. sequence length demonstrates that the predicted mean p(exchange) from Dyna-1 is broad for any given sequence length or helix / strand content (**Fig. 6a**). For proteins at the top and bottom range of mean p(exchange), we discarded proteins as candidates if they were fully intrinsically disordered, did not have published expression/purification methods sections, and already had some relaxation data available. From the remaining proteins, we selected Chitinase 19 from *Bryum coronatum*^*45*^ (BMRB: 11441) with a high degree of p(exchange) (**Fig. 6b**), and hypothetical protein yjbJ^46^ (BMRB entry: 5105) with a low degree of p(exchange) (**Fig. 6c**). Performing ^15^N relaxation dispersion CPMG demonstrated that, resembling Dyna-1’s predictions, the chitinase did have exchange in regions that are crucial for biological activity—in its active site and the loops known to bind chitin (**Fig. 6b,e**)—while the yjbJ only had detectable exchange for residues 2 and 3 (**Fig. 6c,d, supplemental dataset 1**). Evaluated at a per-residue level, Dyna-1 obtained an AUROC of 0.62 for chitinase and an AUROC of 0.72 for yjbJ.

**Fig. 6:**
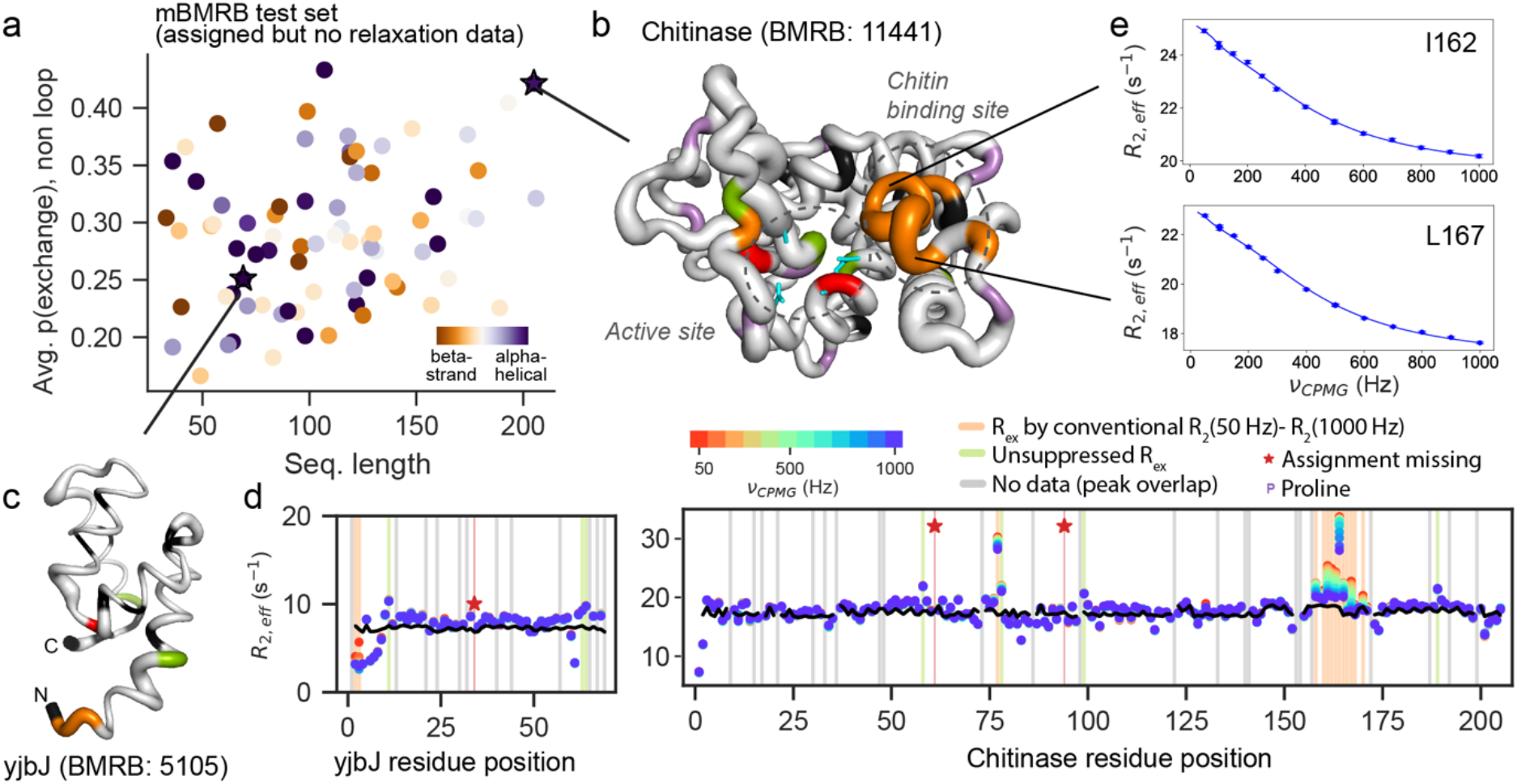
Dyna-1 has predictive power for enzymatic motions in prospective experimental test. a) Dyna-1 proteins in mBMRB test set vary in net p(exchange) predicted across sequence length and helix/strand content. We selected two proteins to experimentally test Dyna-1 predictions: (b) Chitinase 19 from *Bryum coronatum*, and (c) hypothetical protein yjbJ. Residue coloring in (b,c,d) is same as in Fig. 5. (d) ^15^N CPMG relaxation dispersion only showed dispersion curves for two N-terminal residues in yjbJ, whereas chitinase has extensive exchange in the active site and chitin binding site (see also supplemental dataset 1). (e) Representative ^15^N CPMG dispersion data for I162 and L167 in the chitin binding site region.

## Discussion

AF2 fundamentally altered the course of molecular biology and was made possible thanks to the clear task of protein structure prediction defined in the Critical Assessment for Structure Prediction (CASP) challenges and the large amount of standardized data deposited in the PDB. Though widely acknowledged that understanding protein dynamics is essential for understanding protein function, the experimental measurements of dynamics that have uncovered foundational principles of allostery, specificity, catalysis, and many more biological functions have resisted being collected into standardized datasets. Given the predictive power displayed in Dyna-1, it is essential that the field deposits experimental data on dynamics in macromolecules. One barrier to this in NMR has been the perceived diversity and complexity of experiments that measure dynamics. We have laid out in this work a clear task—classify residues with µs-ms exchange—and showed that we could train a model with predictive power across multiple experimental data types: missing assignments, R_ex_ from R_1_/R_2_/hetNOE, and multiple exchange indicators from CPMG.

Dyna-1’s predictive power learned from missing assignments was despite many issues with the current publicly-available data. The data that we used to obtain our labels, lists of atom assignments, are the end point of NMR data processing. The spectra themselves are not required to be deposited, and the lack of standardized requirements to deposit them is a major loss for science. Peak width is related to R_2_^47^, and hence if deposition of even one representative spectrum per assignment dataset had been required, the community could have had roughly the information content of RelaxDB (i.e., R_ex_ labels) for two orders of magnitude more proteins.

Further improvements to Dyna-1 doubtlessly exist for the task of classifying residue dynamics: for instance, ESM-3, the base of Dyna-1, was not trained with sidechain information, information that will likely improve performance. Additionally, Dyna-1 is trained on proteins under 400 residues in length, possibly limiting the scope in which Dyna-1 can predict the dynamics involved in larger proteins. Our comparison of the Dyna-1 model architecture with a training split of 30% sequence identity, 0.5 TM-score cutoff to a larger and more lenient training split of 80% sequence identity, 1.0 TM-score cutoff showed only a marginal increase in performance. This indicates that future model improvements are likely to require innovations in model architectures, rather than increasing data alone.

## Data availability

The mBMRB training set, RelaxDB, and RelaxDB-CPMG datasets are publicly available for noncommercial use at https://github.com/WaymentSteeleLab/Dyna-1 and at https://huggingface.co/spaces/gelnesr/Dyna-1, doi: 10.57967/hf/9277. Raw data for CPMG relaxation dispersion for the two prospective proteins characterized in this study are deposited at 10.5281/zenodo.20834076.

## Code availability

Code to run Dyna-1 is available at https://github.com/WaymentSteeleLab/Dyna-1 and https://huggingface.co/spaces/gelnesr/Dyna-1.

## Author contributions

H.K.W.-S., G.E.N., S.O., and D.K. designed the research. H.K.W.-S. curated the datasets. G.E.N., H.K.W.-S., R.H., H.K. contributed to model training and development. H. K.W.- S. and D.K. interpreted all existing NMR data during model validation. A.M.O. performed all NMR experiments for prospective experimental testing of Dyna-1. H.K.W.-S., G.E.N., D.K. wrote the manuscript. All authors reviewed the manuscript.

## Acknowledgments

We thank Ricardo Padua, Hannes Ludewig, Lucy Colwell, Jeffrey Hoch, Art Palmer, Anthony Gitter, Sameer D’Costa, Kern lab, and Ovchinnikov lab members for useful discussions and advice. We thank Martin Stone for sharing the Indiana Dynamics Database data his group curated in 2000. We thank Katie Henzler-Wildman, Magnus Wolf-Watz, Elan Eisenmesser, J. Patrick Loria, Marcellus Ubbink, George Lisi, Sam Butcher, and Nicolas Doucet for sharing data. We thank Zeming Lin for assistance with ESM-3. Model training was performed on the Stanford Sherlock cluster. We thank Possu Huang for computational and cluster support.

## Supporting Information

## Thorough evaluation of reported NMR relaxation data and corresponding publications increases Dyna-1 performance

While inspecting predictions for other proteins, we noticed that Dyna-1 predicted high p(exchange) for some residues with reported assignments but no relaxation data. This prompted us to revisit our initial curation of experimental data. For instance, in reduced Mercuric Transport protein (MerP)^35^, Thr13 predicts high p(exchange), but has no relaxation data. Ref. ^35^ notes that the peak for Thr13 is very weak, which indicates exchange-broadening and we posit prohibited the authors from fitting R_1_ and R_2_. Though Ala15 was not labeled by our automated labeling scheme, it also has elevated exchange and a large error in R_2_/R_1_ (orange arrow, **Fig. 4c**). Adjusting the experimental label for both residues increases Dyna-1 AUROC from 0.98 to 1.

Another example is the Tudor domain of Survival of Motor Neuron protein (SMNTD)^36^, some of the highest p(exchange) from Dyna-1 were in two residues that were assigned yet had no relaxation data reported: Glu104 and Tyr130 (depicted in black, **Fig. 4c**). The amide hydrogen of Glu104 was assigned at 4.9 ppm, which is a highly unlikely value for an amide hydrogen. We posit that for this small domain and given the well-resolved spectra that the authors report, this residue is more likely exchange-broadened with an incorrect assignment reported. For Tyr130, no R_1_ value was reported, yet the authors categorized it as fitting to “model 4” from the model-free formalism^36^, which is a model containing R_ex_ and requires an R_1_ value. If we consider both residues as having evidence of exchange, Dyna-1’s AUROC increases from 0.67 to 0.75.

In the NtrC receiver domain, Dyna-1 achieved an AUROC of 0.57. We revisited the 10 residues that were assigned yet had no relaxation data. We found that 2/10 had been erroneously assigned in chemical shift depositions corresponding to ref. ^37^ despite being exchange-broadened. 5/10 had very small peaks, indicative of exchange-broadening, and the remaining 3/10 were unanalyzable due to peak overlap. Taking these into account increases Dyna-1’s AUROC to 0.60. Dyna-1 missed predicting exchange in Tyr101 posited to be from ring-flipping. This is exemplary of what we found more broadly across all of RelaxDB, that Dyna-1 is weakest at predicting exchange in beta-sheets (**Fig. 4d**).

Another example is Tyrosine-phosphatase A from *M. tuberculosis* (MptpA)^41^. Dyna-1 has high p(exchange) in loops containing residues with differing orientations between the determined apo and holo structures (apo: 2LUO, holo: 1U2P) and which are postulated to be involved in regulating enzyme specificity^41^. Some of Dyna-1’s high p(exchange) is in the P-loop of MptpA, which our curation indicated had been assigned (BMRB:18533) yet had no relaxation data (BMRB: 26513). However, the authors noted, “Due to exchange broadening the P-loop, residues Cys^11^–Gly^13^ and Ile^15^–Ser^18^ were not detectable”^41^. If we consider these labels, the AUROC for MptpA increases from 0.55 to 0.63.

## Methods

### Curating the “RelaxDB” dataset

Data collected at one magnetic field strength results in three values per residue: a R_1_, R_2_, and hetNOE value. R_ex_ is the increase in R_2_, the backbone amide’s transverse relaxation (see Methods), caused by exchange between states with different chemical shifts on the µs-ms timescale. A common practice to estimate the presence of additional R_ex_ contribution to R_2_ is to look for high outliers in R_2_/R_1_ per residue. R_1_ does not contain R_ex_ and decreases if exchange is present at slower timescales. However, anisotropic tumbling can also alter the R_2_/R_1_ ratio. The model-free formalism^19–21^ was developed to estimate R_ex_ from these data; however, we were unable to robustly apply it at this scale of datasets since relaxation data from multiple field strengths, important for accurate estimations, were often not available. Therefore, we developed the following processing method to label residues with R_ex_. We generated AF2 models for the 163 proteins and calculated their R_2_ and R_1_ values accounting for anisotropic rigid-tumbling only with HYDRONMR^22,23^, fitting one constant per protein that represents the ratio between the experimental and the HYDRONMR overall tumbling time. We then designated a residue as having R_ex_ present if the difference between experimental R_2_ and rigid-tumbling R_2_ (dR_2_) was greater than the combined modelling and experimental uncertainty for that residue (see Methods). We also labeled residues as having ps-ns motion if their hetNOE was less than 0.65 ^24–26^.

#### Theoretical basis for relaxation measurements

The following is condensed from ref. ^1^. The longitudinal (R_1_) and transverse (R_2_) relaxation rates of a backbone nitrogen atom attached to a single proton may be written in a simplified form as

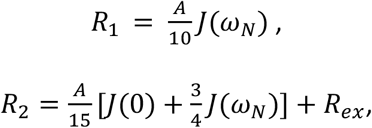

where 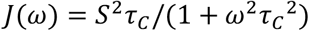. *J*(*ω*) is the spectral density function describing the intensity of exchange occurring at frequency *ω*. These expressions do not include terms on the order of *J*(*ω*_*H*_), which are small on the timescale of protein tumbling (*ω*_*N*_*τ*_*C*_ ≫ 1). In the above, *A* = 3*d*^2^ + 4*c*^2^, where 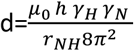 and 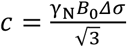. *μ*_0_ is the permeability of free space, *h* is Planck’s constant, γ_*X*_ is the gyromagnetic ratio of atom X (nitrogen N or hydrogen H), *r*_*NH*_is the bond length of the N-H bond, *B*_0_ is the static magnetic field strength, and *Δσ* is the chemical shift anisotropy of nitrogen.

Note that the ratio R_2_/R_1_ for residues that do not demonstrate intramolecular dynamics can be used to estimate the global tumbling time *τ*_*C*_ via the following relationship:

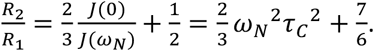

#### RelaxDB curation

We curated 163 R_1_/R_2_/NOE datasets in total. Roughly half of these were deposited in the BMRB, though many required further curation such as converting from incorrectly deposited units. The other half came from supplemental tables of published papers, plot digitization from figures in publications, and private correspondences, including a benchmark of 20 proteins compiled by Goodman et al. in 2000^18^.

##### BMRB entries

To get entries from the BMRB that contained R_1_, R_2_ and hetNOE data, we used https://bmrb.io/software/query/ to download a list of all entries with one entity of polymer type polypeptide(L) that contained R_2_ relaxation data (referred to as T_2_ in the user interface). This resulted in 344 entries when queried in January 2023. We manually went through the list and identified entries that were monomers, had no cofactors present, were not in micelles/bicelles/nanodisks, were not in denaturing conditions, and had at least one set of R_1_/R_2_/hetNOE data all at the same field strength. This resulted in 91 entries. We included data collected at multiple field strengths when available. Several of the BMRB entries had incorrect units, which we fixed by visually inspecting data and comparing to figures of plotted data in the corresponding publications. For three entries, we realized that they contained two or more domains, but that these two or more domains could be modelled as tumbling independently. We therefore separated these into two separate entries.

##### Data from literature

To find data from literature, we searched in PubMed and Google Scholar for keywords “NMR”, “backbone”, and “relaxation”. Upon finding a dataset of an apo, monomeric, stably folded protein and if the data was only available in figure form, we used PlotDigitizer (https://plotdigitizer.com/) to compile values for R_1_/R_2_/hetNOE. We did not include datasets if the figures were too low-resolution to allow for accurately using PlotDigitizer. We only used datasets where we could identify a corresponding set of assignments either cited in the same paper or clearly referencing another set of assignments.

### Fitting data / assigning labels

#### Generating MSAs

We generated MSAs using the ‵run_mmseqs2‵command in ColabFold with ‵filter=True‵. We used these MSAs to predict AF2 models. Prior to calculating conservation metrics (below), we removed sequences with more than 10% gaps and used MMseqs easy-cluster to further filter with options ‵-c 0.9 --cov-mode 1 --min-seq-id 0.9‵.

#### Generating structure models

We used AF2 as implemented in ColabDesign (https://github.com/sokrypton/ColabDesign.git@gamma) to generate the models used for HYDRONMR calculations. We ran AF2 for 3 recycles, seed=0. We used model 1 for each protein. For the Adenylate Kinase datasets (both from *E. coli* and *A. Aquifex*), AF2 predicts the closed state though it is known to occupy the open state predominantly in solution. Therefore, we templated AF2 using structure model PDB ID 4AKE of the open state.

We used these structure models as inputs into HYDRONMR^2,3^ with default settings to estimate R2/R1 from rigid tumbling, denoted as 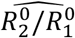. We trimmed unstructured termini of the 161 proteins by removing residues from the N or C termini until the pLDDT of the next residue was over 70, or over 80 for CBFB and PCF11. Residues corresponding to unstructured termini are not considered during evaluation.

#### Fitting R_2_/R_1_ to rigid tumbling estimates

We needed to account for a constant offset in the overall tumbling time between each experimental dataset and the corresponding prediction. To do so, we fit 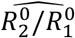 to the experimental data R_2_/R_1_ as follows: for each dataset, we wish to minimize the squared difference between R_2_/R_1_ and 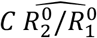 where C is a scalar to be fit, i.e.:

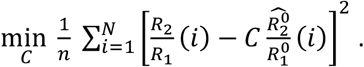

This has a closed-form solution for C:

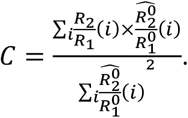

To avoid potential influence of outliers, we first calculate C excluding residues with hetNOE <0.65 as used in refs. ^4–6^ to designate residues with ps-ns motion. We use this C to calculate residuals (i.e., 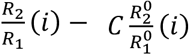, then calculated C again excluding residues with a residual greater than the interquartile range.

Residues exhibiting R_2_/R_1_ greater than that predicted by rigid tumbling, i.e. R_ex_, were identified as those with dR_2_/R_1_ > experimental error + fitting error. Fitting error for each protein was estimated as the interquartile range of the residuals, as this is more robust to outliers than a standard deviation. Experimental error for each residue was taken as the maximum of 5% of the reported R_2_/R_1_ value or the reported error in R_2_/R_1_, whichever value was larger. We also assigned a label for residues with ps-ns motion as residues with hetNOE < 0.65 as used in ref. ^4–6^ to designate residues with ps-ns motion.

All final datasets and corresponding HYDRONMR fits are depicted in **Extended Data Fig. 1**, as well as the normalized RMSE over all R_2_/R_1_ values for residues that were not assigned to have low hetNOE or elevated R_2_/R_1_. RMSE alone would be larger for larger proteins, so we report RMSE that is normalized to the mean R_2_/R_1_ value for each protein. Distributions of norm. RMSE for the final RelaxDB evaluation set, datasets which were initially curated but then excluded, and datasets of proteins removed for containing phosphate buffer and phosphate-related biological function are depicted in **Extended Data Fig. 2a**. In the final dataset, 65/111 datasets had a norm. RMSE of less than 0.1, and 107/111 had a norm. RMSE of less than 0.2 (**Extended Data Fig. 2b**). A norm. RMSE of 0.1 can be interpreted as the experimental data for residues that did not have dynamics assigned were on average within 10% of the predicted HYDRONMR value for R_2_/R_1._

#### Filtering datasets resistant to fitting protocol

After curating 163 datasets initially, we realized in the process of finding a standardized way to fit all of them that some would not be well fit for a variety of reasons. For instance, datasets ISDHN3, MECP2, CBP6C, 5330, and MH35LIF were missing many R_2_ and R_1_ values, datasets 19356, CBP6N, and CBP6C all had hetNOEs < 0.65, and datasets 18773, 6758, and 5762 had relatively high Norm. RMSE values (see **Extended Data Fig. 2a** for comparison of all). We show the raw data for all the excluded datasets in **Extended Data Fig. 1**.

#### Per-residue analysis

For the RelaxDB proteins and the analysis shown in **Fig. 1e**, we calculated conservation scores following the method described in ref. ^7^. To determine buried/exposed residues, we calculated solvent accessible surface area (SASA) using the Shrake-Rupley method^8^ implemented in MDtraj^9^ and normalized values by the maximum value per secondary structure type (loop, helix, sheet) over the whole dataset, following the practice in ref. ^10^. We assigned residues with a normalized SASA ≤ 0.2 as core and > 0.2 as surface residues, following ref. ^10^.

### The “mBMRB” dataset

#### Curating data

We downloaded a list of all entries consisting of polymer type polypeptide(L) from the BMRB^11^ using https://bmrb.io/software/query/. We processed this to retrieve all entities within each entry which were type polypeptide(L). On May 31, 2023, this resulted in 12102 entities. BMRB entries were downloaded using pynmrstar v3.3.1.

We removed entities with isotope labels corresponding to deuterated samples, as a deuterated sample that has not completely back-exchanged with H_2_O would lead to missing assignments. We also removed methyl-labeled samples or samples otherwise partially labeled based on keywords in the isotope metadata. Additionally, we removed any sequences shorter than 30 residues. After this filtering, we had 10123 entities.

We removed any entities with “X” in the sequence (242 removed). Some sequences deposited included multiple copies if the entity consisted of a multimer. We detected and edited these sequences to contain only one copy of the monomeric subunit to match the deposited data. We realized that 1834 sequences contained His-tags, many of which were unassigned. We therefore masked the N- or C-termini of sequences if 6 or more contiguous histidines were present, including everything before the His tag if it was an N-terminal His tag, or everything after if it was a C-terminal His tag. We further filtered the remaining entities to include only entities with 10 or more assignments, to remove entries that included only a few backbone chemical shifts (removed 273).

For each residue, we assigned labels corresponding to whether the ^15^N backbone chemical shift is missing or not based on if that residue had a chemical shift entry corresponding to an “Atom_ID” of N.

We further removed entities that had 15 or more contiguous missing assignment labels between any assigned residues. This was to filter entities where we noticed that entire domains were unassigned. We selected the cutoff of 15 contiguous residues because the most residues missing in a row in the RelaxDB dataset was 14. This removed 227 entities.

Of the remaining 9,381 entries, 5% did not have isotope information present. However, we elected to keep these, as we saw that they had on average fewer missing assignments and shorter sequences than the 95% of other remaining sequences which had 1H labeling (**Extended Data Figure 3a,b**). Roughly 60% of these are older entries (entry_ID < 10000).

**Extended Data Fig. 3d** contains average fraction missing across the mBMRB divided by pH, temperature, and sequence length. From this we observe that there are certain well-known factors that play a role in more missing assignments—for instance, pH greater than 8, or sequence length greater than 250—but these entries also represent small fractions of the data in total. Understanding how to more systematically account for such systematic outliers will be the topic of future research.

Structure models for each protein in the mBMRB dataset were generated using ESMFold v1^12^.

#### Comparing to structure models in PDB

We used MMseqs^13^ to align all the mBMRB sequences (9,381), filtered to 80% sequence identity, against all protein chains in the PDB (curated based on PDB data in NMRbox, October 2024). We kept all hits with a bitscore greater than 50 and sequence identity greater than 90%. We kept one chain per protein sequence per model. We found that 95% of the mBMRB had one or more similar PDB structures.

### RelaxDB-CPMG curation

#### Complete dispersion data available

AdK: R_2,eff_ vs ν_CPMG_ for apo AdK from *Aquifex aeolicus* at 10°C collected at 800 MHz, presented in ref. ^14^, as well as a corresponding peak list, was found in the Kern lab data archive, with thanks to K. Henzler-Wildman.

BlaC: R_2,eff_ vs ν_CPMG_ of apo BlaC collected at 600 MHz and 850 MHz, presented in ref. ^15^, was kindly provided by M. Ubbink. A peak list to identify missing assignments was taken from BMRB entry 27888 (same as in RelaxDB).

BLVRB: R_2,eff_ vs ν_CPMG_ of apo BLVRB, presented in ref. ^16^, was kindly provided by E. Eisenmesser. A peak list to identify missing assignments was taken from BMRB entry 27462.

CypA: R_2,eff_ vs ν_CPMG_ of apo CypA, collected at 10°C and 25°C at 600 MHz in ref. ^17^, as well as peak lists to identify missing assignments at both temperatures, were found in the Kern lab data archive. Because different processes are more prominent at 10C° or 25C°, the final labels created represent whether a residue experienced exchange at either temperature.

VHR: R_2,eff_ vs ν_CPMG_ was taken from 800 MHz data provided in Table 7.10.3 of ref. ^18^. Missing assignments were taken from assignments in Table 7.10. Note that these differ from assignments by the authors in BMRB 27950; the assignments in the dissertation table are more complete.

K-Ras: R_2,eff_ vs ν_CPMG_ of K-Ras with GTP bound, presented in ref. ^19^, had been publicly deposited at https://doi.org/10.5061/dryad.j6q573nm0.

#### Complete dispersion data unavailable

CheY: Residues with missing assignments were taken as the black residues in Fig. 2 from ref. ^20^. Residues were labeled as having exchange when CPMG could individually be fit (all residues in Table 2 in ref. ^20^). This is CheY in the presence of 1 mM EDTA to remove any Mg^2+^, which alters the dynamics.

HtrA2 PDZ domain: Values for R_ex_ were taken from source data corresponding to Fig. 2a in ref. ^21^. Assignments were taken from BMRB:51320.

RNase: Residues were labeled as having exchange when R_ex_ >3 Hz at 20°C in either 500MHz or 600 MHz data. This recreates Fig. 4B in ref. ^22^. The R_2,eff_ vs ν_CPMG_ data for these residues was kindly provided by P. Loria, but the dispersion data for all residues was not available.

Arginine kinase: Labels for residues with exchange were taken from Table 1 in ref. ^23^, which summarized all residues for which CPMG data could be fit individually. Table S1 in ref. ^23^ indicated if residues were missing due to exchange-broadening or overlap, and we used those labels.

### Identifying unsuppressed R_ex_ in CPMG experiments

We first compared model predictions to labels for exchange where we assigned labels using a conventional assignment from the literature: any residue where R_ex_ is significantly apparent, calculated as R_2,eff_ (first field strength) - R_2,eff_ (last field strength) is greater than background noise.

However, we noticed that in some datasets there were residues with noticeably high R_2,eff_’s across all field strengths but R_2,eff_(first) - R_2,eff_(last) was not substantial enough to be greater than 2-3 Hz. These correspond to residues that have exchange at the microsecond timescale: at the fast end of the micro-millisecond regime that CPMG can detect.

However, assigning these residues with confidence requires determining what residues have elevated R_2,eff_ when there is no suppression, the equivalent of measuring R_ex_ as is conventionally done. Given our success using HYDRONMR to quantitatively fit R_2_/R_1_ ratios in RelaxDB, we used HYDRONMR to calculate R_2_values and fit a scalar to find the closest fit between R_2,inf_ and R_2,rigid_.

The R_2,eff_ data for all residues, corresponding R_2,rigid_ values, and labels are in **Extended Data Fig. 8**.

### Data splitting

We clustered sequences using MMseqs^13^ ‵easy-cluster‵ with arguments ‵--cluster-mode 1 --min-seq-id 0.8 -k 5‵. We used the resulting clusters to remove all sequences similar to those in RelaxDB or RelaxDB-CPMG. From the remaining sequences, we randomly separated out 500 clusters for the validation set and 100 clusters for the test set.

For training with a 50% or 30% sequence identity cutoff, we used –min-seq-id 0.5 [or 0.3] to cluster but kept the same proteins in the validation and test set. To create training splits with structure homologues removed at TM-score = 0.5 or 0.7, we calculated all pairwise TM-scores using TMalign^24^ between the mBMRB-Train set and mBMRB-Val, mBMRB-Test, RelaxDB, and RelaxDB-CPMG sets. We used TM-score normalized by the average length of the query and target protein, as recommended in ref. ^24^.

The model sweep depicted in **Fig. 3c** was performed with a different dataset split that considered all 163 sequences initially curated for RelaxDB and used a split of 80% sequence identity, 1.0 TM-score cutoff. The splits used for the model sweep in **Fig. 3c** can be found in the ‵oct2024‵ data repository. Subsequent model training was reperformed using a data split considering only the later filtered 133 sequences in RelaxDB and used the most stringent training data split (30% sequence identity, 0.5 TM-score cutoff). These subsequent models are depicted in **Figs. 3d** and onwards.

### Baseline training

Our baseline training aimed to test simpler representations of protein sequences and/or structure. We use the mBMRB data to train these methods. The results of these baseline models on the mBMRB-Validation, mBMRB-Test, and RelaxDB datasets are depicted in **Extended Data Fig. 4a**. None outperform the models based on pre-trained models.

#### Naïve baselines

The simple, naïve baseline uses amino acid frequencies in the mBMRB training data as weighting of a random classifier.

- The simplest is the AA baseline, which computes the frequency of each amino acid being missing from the mBMRB training data. We then predict for our validation and test set whether a residue is missing at random, weighted by that relative frequency.
- We evaluate similar random classifier using frequency of secondary structure (loop, sheet, helix) as calculated by DSSP, and a combination termed the “AA & DSSP” baseline using the frequency of each amino acid type given its secondary structures (loop, sheet, helix).

#### Simple classifiers

We use a similar Transformer classifier as that described for the final Dyna-1 architecture trained on simpler representations of the input mBMRB training data. Each representation is described.

- “OHE AA”: we input each protein as a one-hot encoded sequence, where each residue is represented by a unique combination of 0s and 1s, into the transformer classifier.
- “SASA” we evaluate solvent-accessible surface area using the Shrake-Rupley method^8^ implemented in MDTraj^9^. These values are encoded by a simple one-dimensional MLP of hidden size 128 and then input into the transformer classifier.
- The “SASA & DSSP”, “SASA & AA”, and “SASA & AA & DSSP” concatenate the outputs of the MLP with a one-hot encoding of the secondary structure, sequence, and secondary structure and sequence, respectively. This concatenated embedding is subsequently used as input into a transformer and trained accordingly.

#### Other representations

- Normal mode analysis (NMA) is a computational technique that has long been of interest for modelling dynamic processes in proteins^25^. To evaluate NMA, we use ProDy^26^ to compute the top 20 normal modes of the models generated using ESMFold (for the mBMRB data) or AF2 (for the
- RelaxDB data) using the Anisotropic Network Model (ANM) and Gaussian Network Model (GNM) and build the Hessian matrix and Kirchhoff matrix, respectively. We then calculate the mean square fluctuation and evaluate the AUROC and AUPRC using these values. No model is trained.
- To use sequence conservation as a baseline, we computed sequence conservation scores and coverage index per protein as described in ref. ^7^ and evaluated the AUROC and AUPRC using these values as logits. No model is trained.

### Deep learning training

We trained Dyna-1 on the mBMRB-Train dataset, sampling using a sequence-cluster-based approach. One protein per sequence cluster is chosen randomly during each epoch. We used the validation set to compute the validation loss and validation metrics. The final model weights for each training run for all models are chosen using max AUROC on the validation set. We excluded unassigned termini and prolines (which have no backbone amide and therefore are always missing) from model evaluations.

We used the ESM2-t30-150M model for ESM-2 embeddings, which has 31 layers and 150M parameters. Initial attempts using ESM2-t6-8M and ESM2-t12-35M indicated that smaller models did not perform as well, and ESM2-t33-650M performed marginally similar. We used the outputs of AF2 model 1 to construct the AF-pair representation. The weights from ESM and AF2 were fixed (frozen) during training.

Models were trained with the AdamW optimizer^27^ with a learning rate of 1e-6. The AF2-pair model was trained with batch size of 4 with an accumulation step every 4 epochs, while the ESM-2 and ESM-3 based models were trained with a batch size of 16 and accumulation each epoch. Training is completed within 500 epochs, and we use early stopping when validation loss begins to overfit. We choose our best performing model by comparing normalized AUPRC on the validation set, as we saw this had slightly more discrimination than AUROC. Normalized AUPRC is calculated by subtracting the baseline AUPRC from the AUPRC for each protein.

The transformer architecture used for the ESM-2 embeddings has 8 heads and 12 layers. The transformer for the ESM-3 embeddings has 6 heads and 12 layers, and the transformer for AF2-pair representation has 4 heads and 4 layers. We note that increase in number of heads or number of layers had negligible improvements for ESM-based models, and we were limited by GPU memory for AF2-pair. The hidden dimension for the ESM-based models is equivalent to the hidden size of the embedding, while for the AF2-pair model is 128. We apply a 0.1 dropout rate. All models were trained on a single NVIDIA A40 GPU, which takes between 12 hours and 2 days to complete training.

### Adjusting logits to account for class imbalance

When converting from logits to probabilities, we adjusted the raw logit outputs predicted by the model to account for the significant class imbalance in the problem following the practice in ref. ^28^ (cf. **Supporting Information Fig. 1**). This practice seeks probabilities that minimizes the “balanced error”, the prediction error averaged over all classes. In brief, for data *x* labelled with classes *y*, the native posterior probability *P(y* | *x)* is proportional to the prior probability of class y, *π*(*y*):

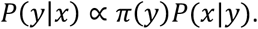

In seeking to minimize the balanced error, the balanced class-probability function instead implicitly becomes

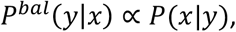

where *L* is the number of classes (L=2 in our binary classification context).

Menon et al. describe how *P*^*bal*^(*y*|*x*) is a Bayes-optimal scorer for minimizing the balanced error. For a Bayes-optimal scorer, the accuracy of any given class is not proportional to its degree of representation in the training data, which is desirable in our context.

Consider our model to be a scoring function *ŝ*, with learned class probabilities 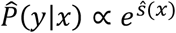. Via the definition of *P*^*bal*^, we can approximate the Bayes-optimal scorer using our given scorer by using an adjusted scorer *s*′:

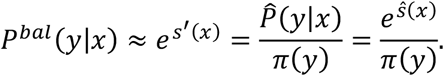

Accordingly, the adjusted logits may be written as *s*^′^(*x*) = *s*^′^(*x*) − log *π*(*y*).

In the context of our binary classification problem, let *π*^*m*^ be the fraction missing in the training data set and let *l(x)* be the raw logit output given input *x*. Dyna-1 is structured to just predict one logit, the logit of the “missing” class, hence the logit predicted for the “not missing” class is always zero. We calculate our probability P(missing) as

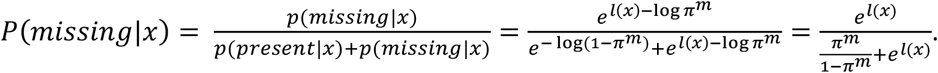

This can be rewritten in terms of the sigmoid function *σ*(*x*) = (1 + *e*^−*x*^)^−1^, which is typically used to convert logits to probabilities:

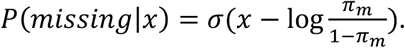

**Supporting Information Figure 1.**
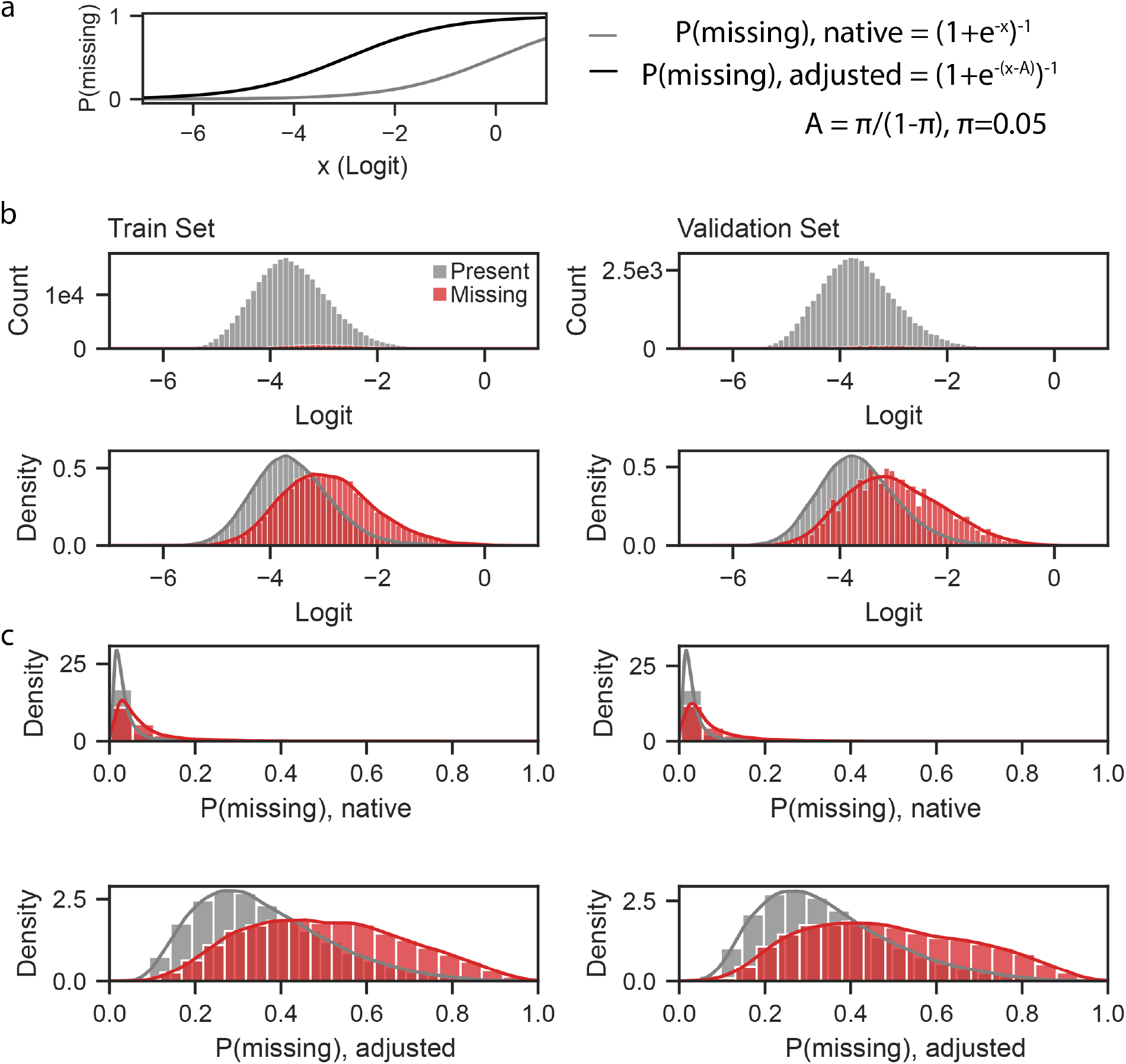
Adjusting native logits from Dyna-1 to account for class imbalance. (a) We adjusted probabilities calculated from logits to maximize the “balanced accuracy”, i.e. accuracy averaged across both classes, rather than accuracy averaged over all datapoints (see Methods). We set our prior for the missing class based on the training data, π, as 5%. Comparing the adjusted curve (black) to the probability densities in (b) demonstrates how this intuitively matches where a discriminator between the two classes should be (at x ~ −3). (b) Histograms and probability densities of logits predicted for residues labeled as “present” and “missing” over train and validation sets. (c) Densities of resulting p(missing) if calculated using native logits or adjusted logits.

### Uncertainty estimation

We estimated uncertainty for comparing models in the RelaxDB dataset (**Fig. 4a, Extended Data Fig. 4**) using a 95% confidence interval calculated in Seaborn v0.11.2 ^29^.

We estimated uncertainty in binned residue analyses (**Fig. 4e,f**) using bootstrapping. We resampled residue subsets with replacement to obtain 100 bootstrap iterations that contained at least one of each class (required for calculating AUROC).

All depicted p-values (**Fig. 1e, Fig. 2h**) are for a 2-sided hypothesis test whose null hypothesis is that two sets of data are uncorrelated, performed using statannotations v0.2.3 (https://pypi.org/project/statannotations/).

We curated uncertainties for the RelaxDB datasets in the following way: if authors reported R_1_/R_2_/hetNOE uncertainties, we used those. In the BMRB, we found a few with a deposited uncertainty of zero. We set these to be the smallest nonzero uncertainty value given for that data type in the dataset. If no uncertainties were given, we set the uncertainty for R_1_ as 10% the measured value, and 5% for R_2_ and hetNOEs. If the data was plot digitized and had error bars, we digitized the error bars as uncertainties.

### Preparation of recombinant proteins

The hypothetical protein yjbJ (YjbJ_ecoli) (NCBI: WP_058905752.1) and wild-type *Bryum coronatum (*BcChi-A) constructs were ordered from GenScript (Supplementary Table 1). The codon-optimized YjbJ_ecoli sequence was cloned into a pET-15b expression vector with an N-terminal 6xHis tag followed by a TEV protease cleavage site. The wild-type BcChi-A (without any affinity tag) was cloned into a pET-22b vector between the Ndel (5’) and BamHI (3’) restriction sites, replacing the pelB leader sequence of the vector.

The individual plasmids encoding each protein were transformed into *E. coli BL21(DE3)* cells (New England Biolabs). Uniformly ^13^C, ^15^N-labeled proteins were expressed in *E. coli* grown in 1L M9 minimal medium supplemented with 2.0 g/L U-^13^C as the main carbon source and 1.0 g/L of ^15^NH_4_Cl (Cambridge Isotope Laboratories). Cells were grown to an OD_600nm_ of 0.6 before induction with 1 M isopropyl thiogalactoside (IPTG). Then grown overnight at 25°C for the BcChi-A construct or 15 °C for the YjbJ construct for ~18 h.

Both constructs were purified using a similar method as previously described ^30,31^. Briefly, YjbJ cell pellets were resuspended in 20 mM Tris, pH 8, 500mM NaCl, 30mM Imidazole (Buffer A), DNAse 1 (Sigma-Aldrich), and lysozyme (Sigma-Aldrich). Lysate was sonicated on ice for 10 min (20 sec on, 30 sec off, output power of 40 W), followed by clarification via centrifugation at 19000 rpm for 45 min at 4°C. A 5 ml His-trap Ni^2+^ affinity column (Cytiva) was pre-equilibrated in buffer A. The column was washed with 10 column volumes of buffer A after loading the supernatant, followed by elution of the his-tagged YjbJ protein with buffer containing 500 mM imidazole. The His tag was cleaved off with TEV protease and removed by loading the cleavage reaction back onto the His-trap Ni^2+^ affinity column. YjbJ with no tag was collected in the flow-through with 5 column volumes of buffer A containing no Imidazole. The ^13^C,^15^N labeled YjbJ was further purified on a Superdex 75 26/600 pg column in 25 mM Na_2_PO4, pH 6.5. BcChi-A construct was purified by resuspending the pellet in 20mM Tris-HCl, pH 7.5, and sonicated on ice. The lysate was spun down at 18500 rpm for 45 min, followed by dialysis of the supernatant against 10 mM sodium acetate, pH 4 and additional clarification by spinning down at 18500 rpm for 20 min. The supernatant was then dialyzed in 10 mM sodium acetate, pH 5, before loading on a 5 ml HiTrap Q HP column (Cytiva) pre-equilibrated in the same dialysis buffer. The protein was eluted with a linear gradient of NaCl from 0 to 150 mM NaCl in the same buffer. The ^13^C, ^15^N labeled BcChi-A was further purified on a Superdex 75 26/600 pg column in 50 mM Acetate, pH 5.

### NMR spectroscopy

^15^N-Carr-Purcell-Meiboom-Gill (CPMG) data were carried out on a 900 MHz Bruker spectrometer equipped with a cryogenic triple resonance probe. NMR data were collected and processed using TopSpin 3.2. Measurements for BcChi-A were performed at 300 K on a 1 mM ^13^C, ^15^N labelled sample in 50 mM Acetate, pH 5.0, while YjbJ experiments were conducted at 298 K on a 1 mM sample of ^13^C, ^15^N labelled in 25 mM Na_2_PO_4_, pH 6.5. ^15^N-CPMG^32,33^ data was acquired as pseudo 3Ds with a fixed total CPMG relaxation period (T_CPMG_) of 40 ms, and 16 *n*_*cyc*_ values ranging from 1 to 40. A reference spectrum was collected with no CPMG relaxation delay (*n*_*cyc*_=0), and duplicate measurements at *n*_*cyc*_ of 4 and 20 were collected for error estimation. The effective transverse relaxation rate *R*_*2,eff*_ was calculated from these datasets, and errors were taken as the larger of noise-based versus duplicate-based error estimation. Each dataset was collected with 32 scans per FID for the BcChi-A or 8 scans per FID for the YjbJ protein with 2 sec repetitive delay between scans. All CPMG traces for both proteins are in Supplemental Data 1.

Backbone assignments for the BcChi-A and YjbJ were obtained from BMRB accession codes 11441 and 5105, respectively.

### NMR data analysis and fitting

Peak heights for individual residues were collected from each 2D ^15^N-HSQC in the pseudo 3D ^15^N-CPMG by lineshape fitting using PINT^34^ software. The R_2,eff_ at a given CPMG field strength (v_cpmg_) was calculated using:

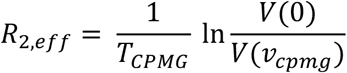

where T_cpmg_ is the constant CPMG relaxation period, V(0) is the peak volume and V(v_cpmg_) is the peak volume at each CPMG field strength.

Relaxation dispersion curves, expressed as *R*_*2, eff*_ versus CPMG field strength, were fit per residue using ChemEx software (https://github.com/gbouvignies/chemex) ^35^.

## Extended Data

**Extended Data Figure 1:**
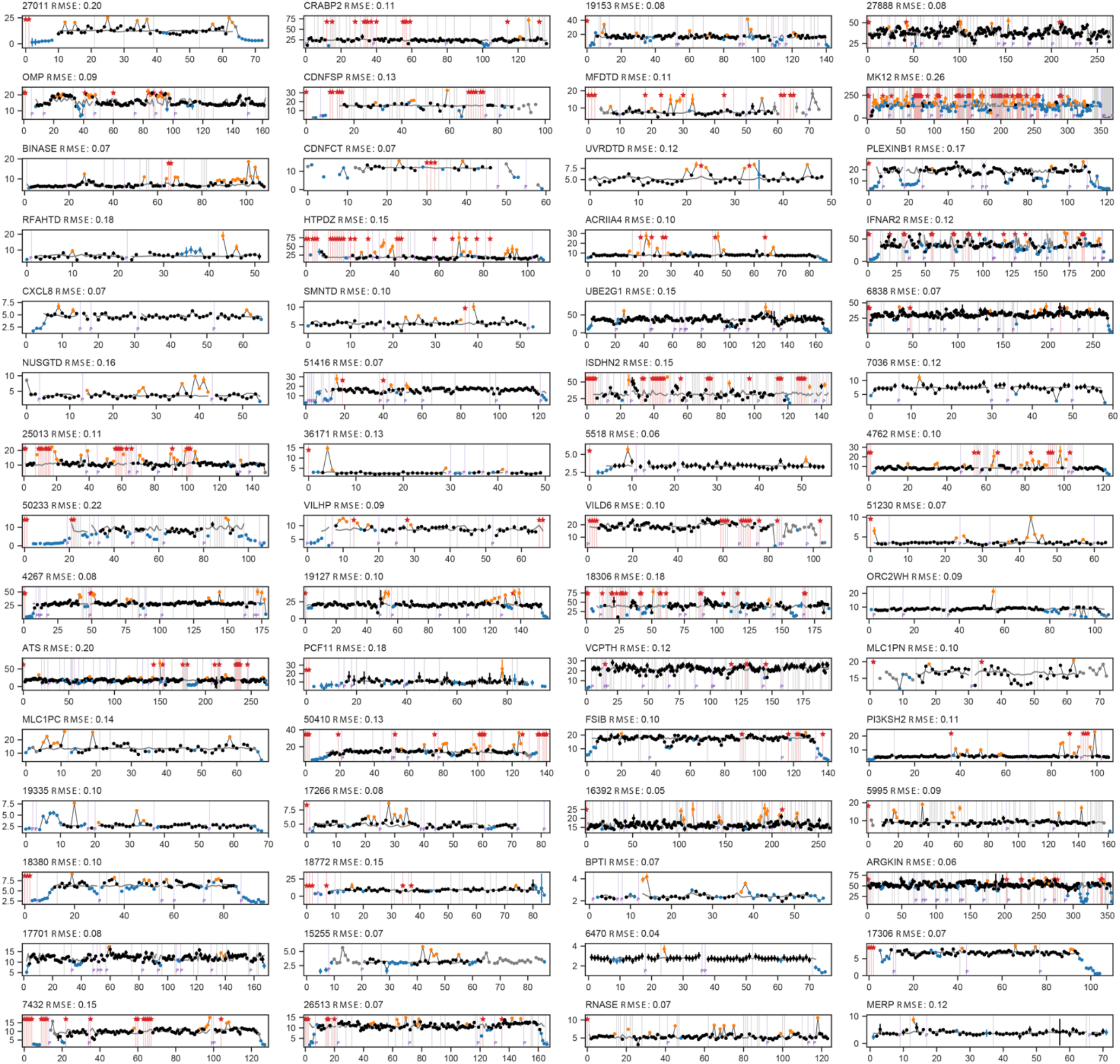
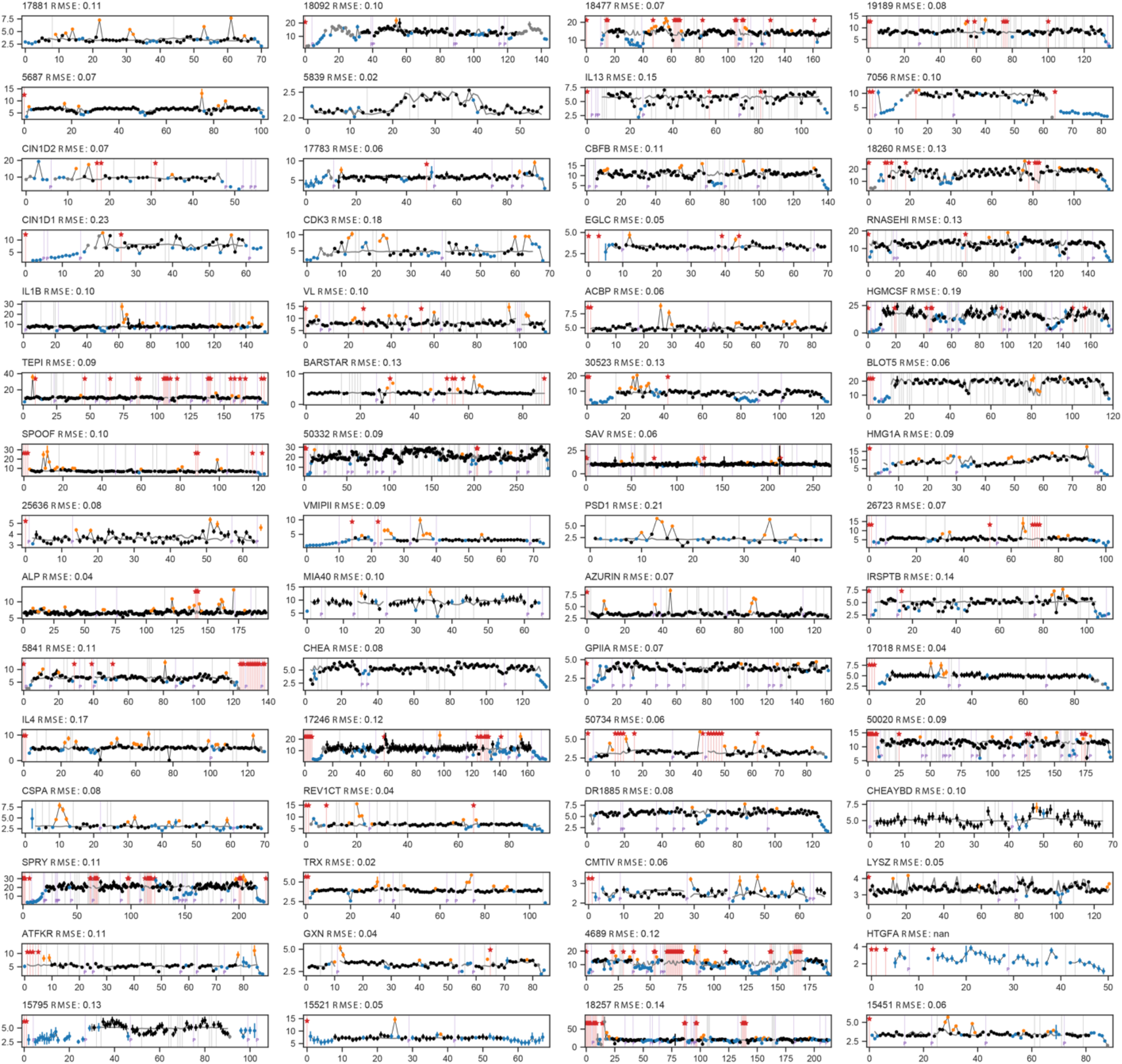
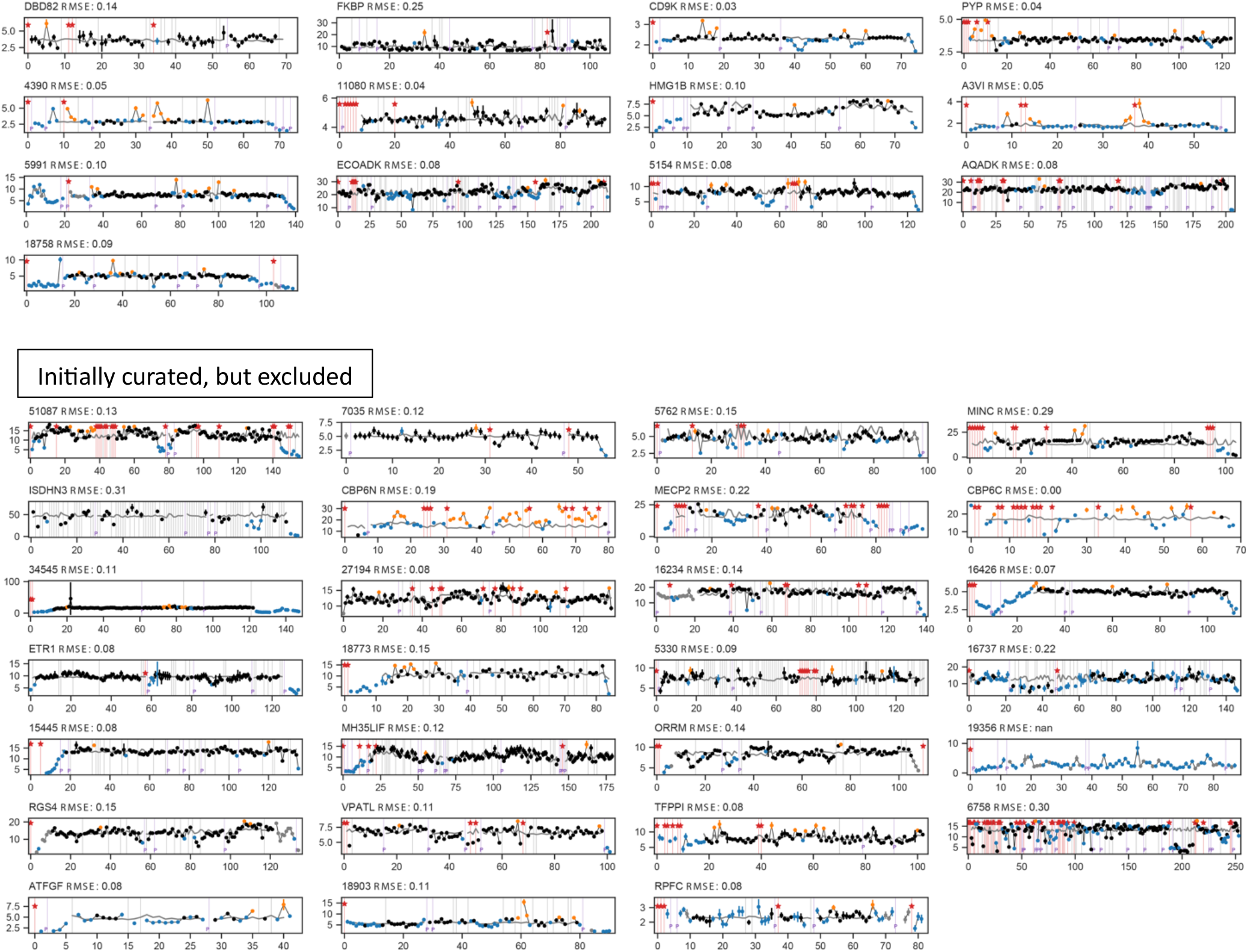
(page 1/3) ^15^N R_2_/R_1_ data and labels for all 133 proteins in RelaxDB and the 30 which were initially curated but excluded from final evaluation set for various reasons (see Methods). Calculated R_2_/R_1_ from HYDRONMR^25,26^ for rigid tumbling is depicted by grey line. Colors used are the same as in Fig. 1. Title for each is RelaxDB entry ID followed by normalized RMSD.

**Extended Data Figure 2:**
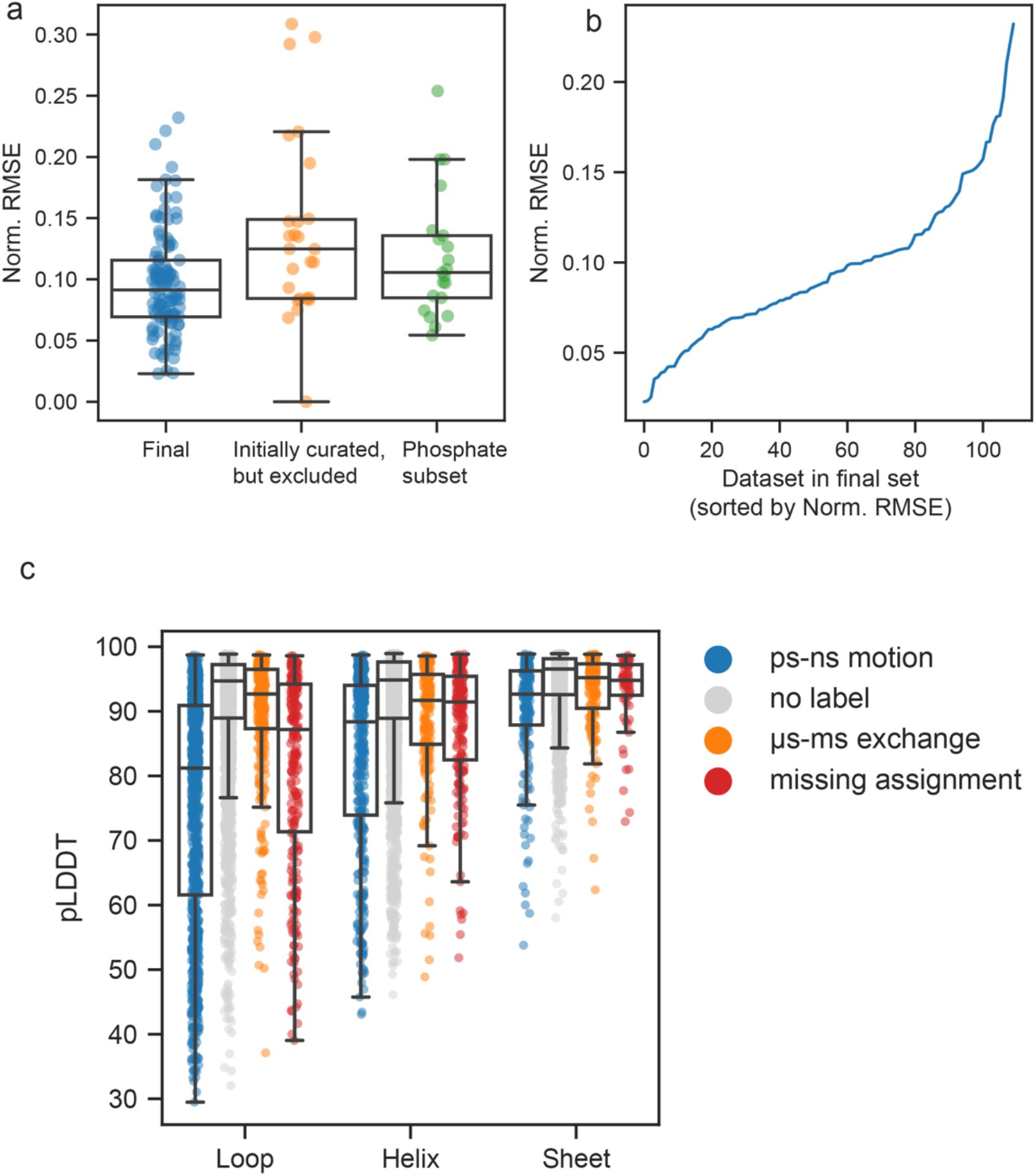
(a) RMSE normalized to median R_2_/R_1_ per protein for the final RelaxDB dataset, protein datasets that were initially curated but excluded, and proteins with biological function involving phosphate binding, but measured in phosphate buffer. n=160 proteins. (b) Norm. RMSE distribution of final RelaxDB dataset used for evaluation. (c) pLDDT (predicted local distance difference test) from AF2 by secondary structure and dynamics type. Low AF2 pLDDT is most indicative of ps-ns motion in RelaxDB. Box plots depict median and 25/75% interquartile range, whiskers = 1.5 *interquartile range, n=12,584 residues.

**Extended Data Figure 3:**
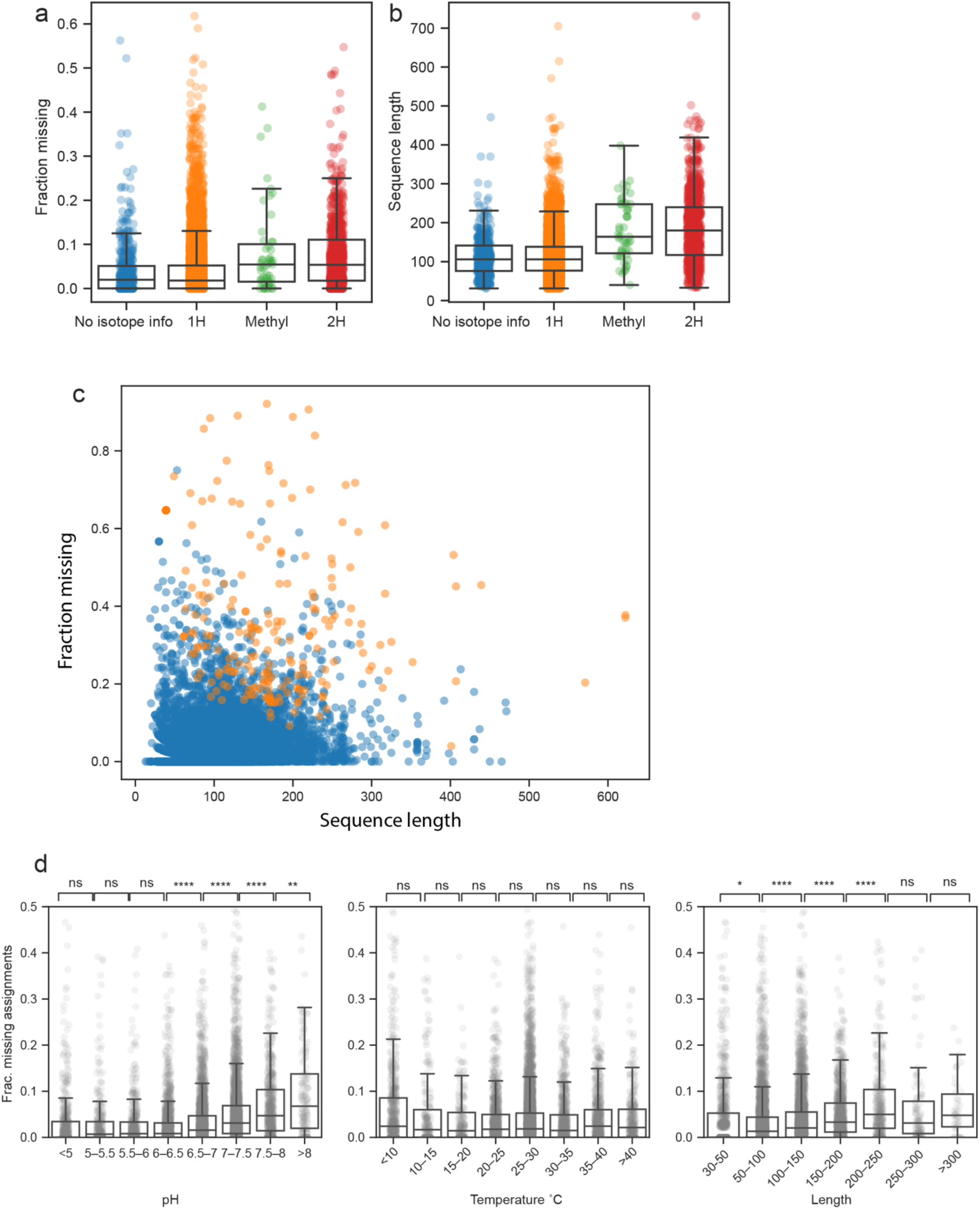
Statistics from curating the “missing assignment BMRB” (mBMRB). We removed proteins with deuteration or methyl or specific labelling, based on metadata. Some proteins had no isotope information, yet we elected to keep them, as they had fewer missing assignments compared to proteins with ^1^H isotope labels and were overall shorter than other isotope categories. (a) Fraction missing split by isotope category (n=11,268 proteins). (b) Sequence length by isotope category (n=11,268 proteins). (c) We removed entries with 15 or more consecutive missing assignments (orange). (d) Missing assignments as a function of pH, Temperature, and sequence length. In (a,b,d): Box plots depict median and 25/75% interquartile range, whiskers = 1.5 times the interquartile range. In d: statistical comparisons by two-tailed independent-samples t-test with multiple comparisons adjustment. ns: 0.05 < p <= 1. *: 0.01 < p <= 0.05. **: 0.001 < p <= 0.01. ***: 0.0001 < p <= 0.001. ****: p <= 0.0001.

**Extended Data Figure 4.**
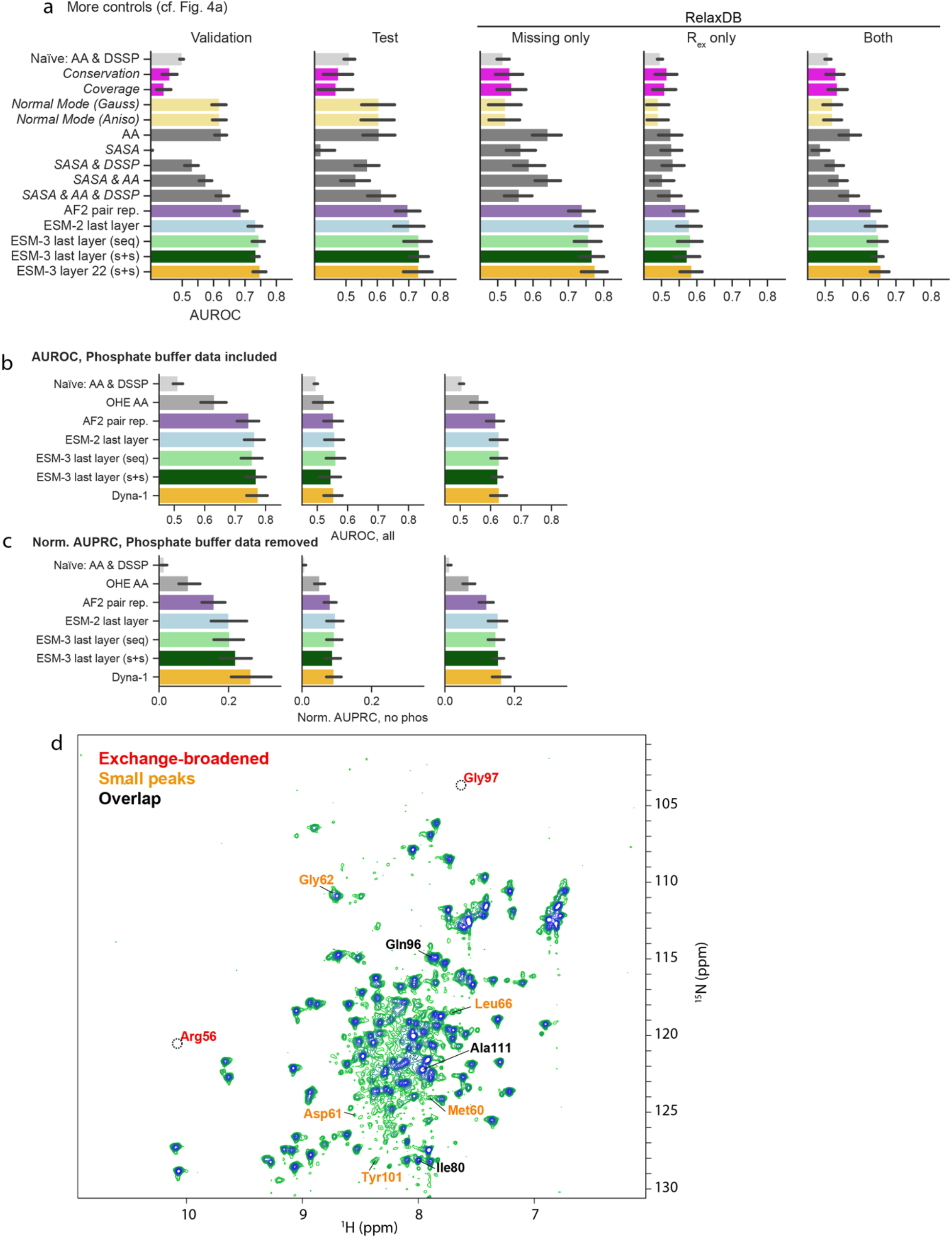
(a) Validation, test, and RelaxDB datasets comparing further benchmarks at training cutoff of 30% sequence identity, TMscore 0.5. Benchmark models in italics are not in Fig. 4a. (validation: n=500 proteins, test: n=100 proteins, RelaxDB: n=112 proteins). (b) RelaxDB evaluated by AUROC across different models tested, using all 133 proteins including those with exchange from phosphate binding (cf. **Fig. 4a**). (c) RelaxDB evaluated across different models evaluated by normalized AUPRC, with phosphate binding proteins removed (n=112 proteins). In (a,b,c), bars represent mean, error bars represent 95% confidence interval evaluated over proteins in RelaxDB. (d) ^1^H-^15^N HSQC of NtrC at 25°C, annotated with residues for which assignments exist in BMRB entry 4527 (non-phosphorylated NtrC), but relaxation data is missing from BMRB entry 4762. The Arg56 and Gly97 assignments in entry 4762 (shown in red here) are presumed to be erroneously copied from entry 4528, which are assignments corresponding to phosphorylated NtrC. All residues labeled orange are very weak peaks indicative of R_ex_. For these, we concluded exchange broadening was the reason for no reported relaxation data. Black residues are overlapped peaks.

**Extended Data Figure 5:**
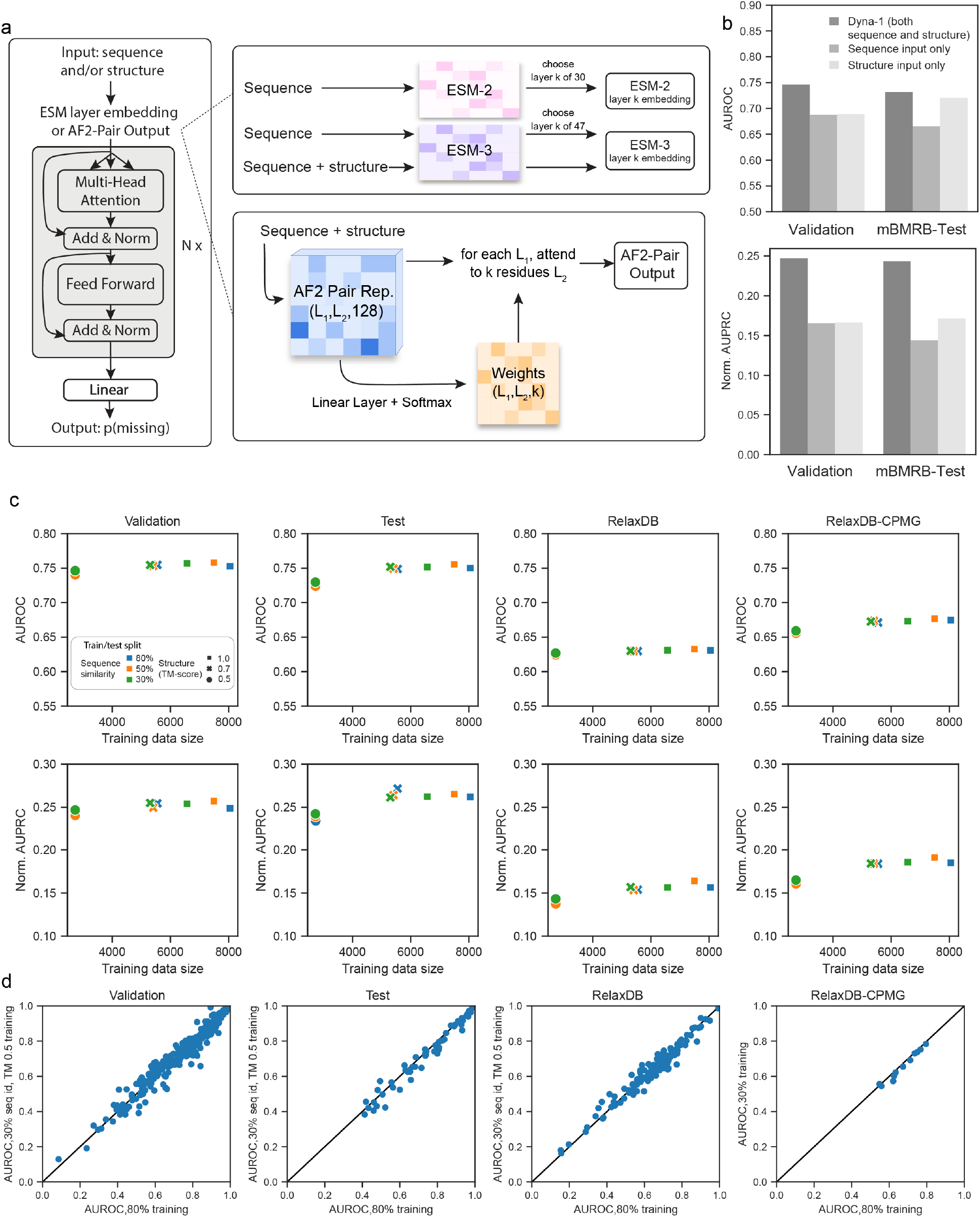
Details of training Dyna-1. (a) Architectures of tested ESM and AF2 inputs, outlining the model architecture. At training time, either a sequence-structure pair is passed through AF2 or ESM-3, or a sequence is passed through ESM-2 or ESM-3. The corresponding embedding from the k^th^ layer from ESM or AF2-pair output is fixed (“frozen”) and used as input into a one-dimensional Transformer architecture. Output is one value per residue, the logit for that residue being missing. (b) Comparing Dyna-1 performance given both sequence and structure (blue) vs. sequence only (orange) or structure only (green). Performance is best when given both sequence and structure. (c) Dyna-1 performance given different training split cutoffs based on sequence identity and structure similarity (cf. Fig. 3d) across all datasets. (d) AUROC per protein for either training with the 80% sequence identity and 1.0 TM-score split or 30% sequence identity and 0.5 TM-score split. Performance on individual proteins in the validation, test, RelaxDB, and RelaxDB-CPMG does not significantly change between more stringent sequence similarity cutoffs.

**Extended Data Figure 6.**
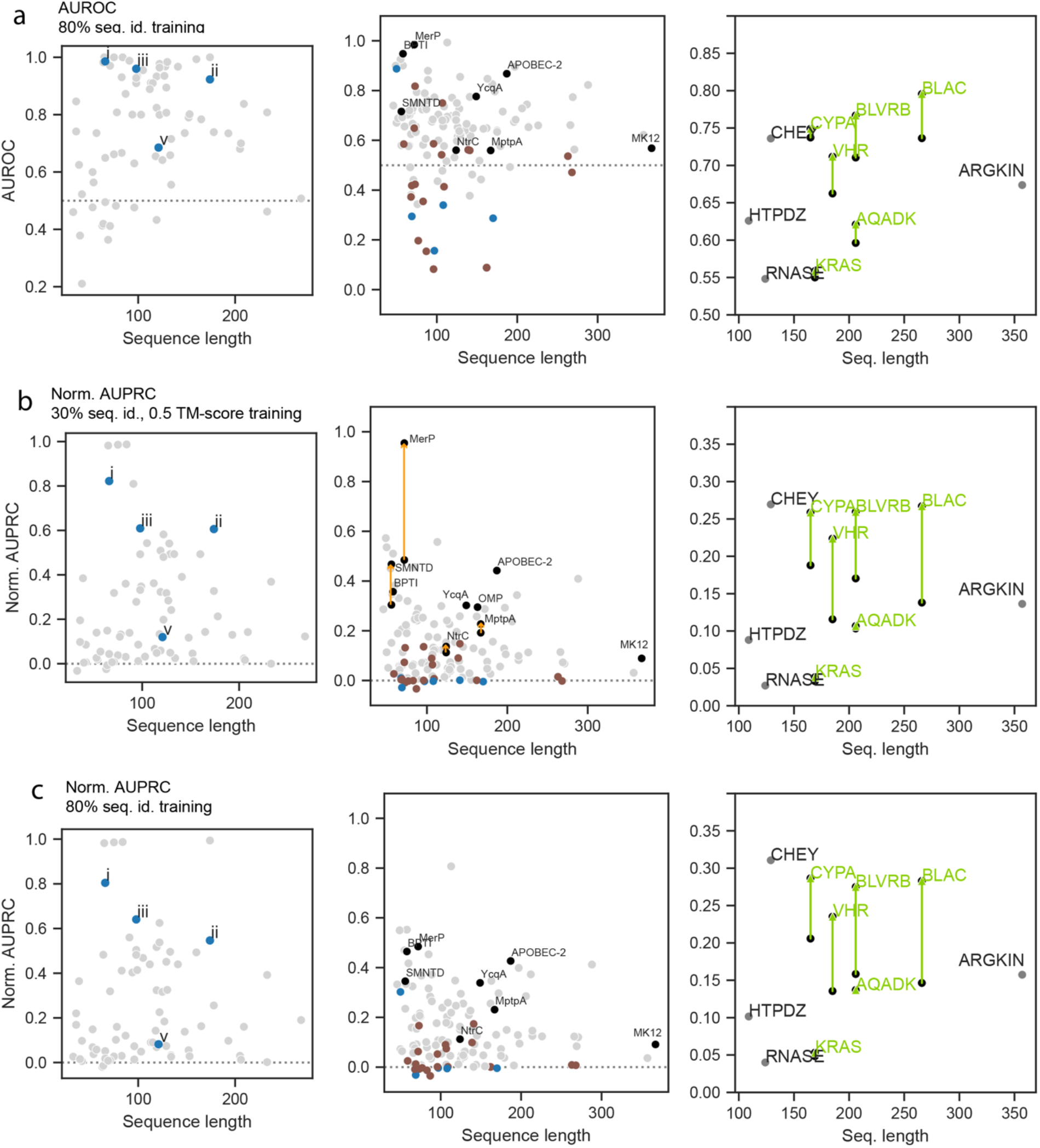
Dyna-1 trained with 80% sequence identity cutoff is comparable to results presented in main text for 30% sequence identity and 0.5 TM-score cutoff. (a) AUROC for mBMRB-Test (cf. Fig. 3e), RelaxDB (cf. Fig. 4b), RelaxDB-CPMG (cf. Fig. 5d). Coloring for each follows respective main-text figures. (b) Normalized AUPRC per protein for Dyna-1, trained with 30% sequence identity and 0.5 TM-score cutoff. (c) Normalized AUPRC per protein for Dyna-1, trained with 80% sequence identity cutoff.

**Extended Data Figure 7:**
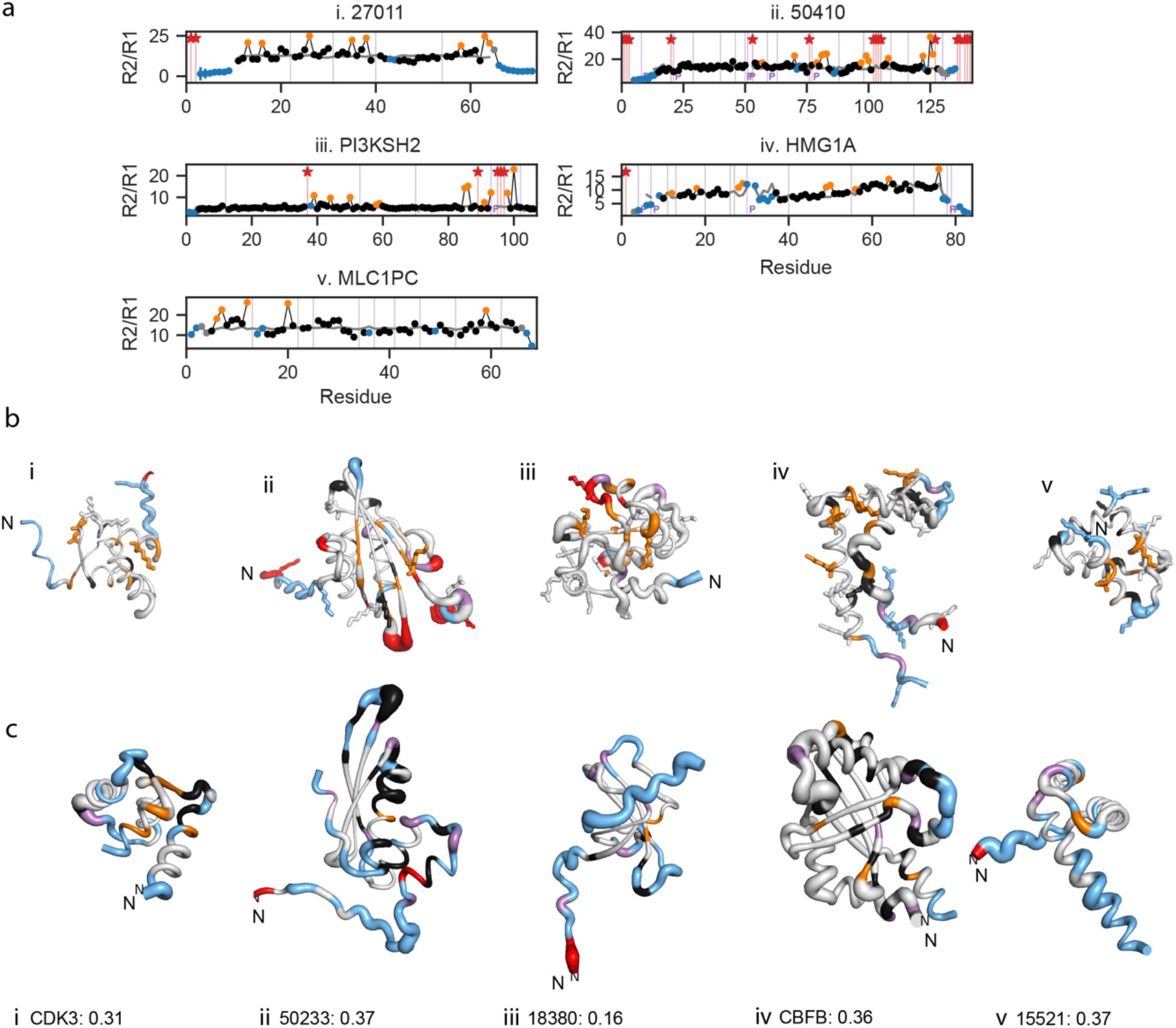
Some proteins with poorly predicted exchange in RelaxDB reflect phosphate-binding exchange or high probability for ps-ns motion. Numerous proteins in RelaxDB predicted with an AUROC < 0.5 are these proteins that bind phosphate ion moieties during their biological function, yet the NMR experiments were measured in phosphate buffer. (a) R_2_/R_1_ data for 6 of these proteins which bind RNA or DNA. (b) All positively charged sidechains (arginine, lysine) are depicted in stick form. Many of the residues with experimental exchange (orange) that are not predicted by Dyna-1 are on the surface of these proteins and can be explained by phosphate binding. (c) Proteins with low AUROC and high p(exchange) predicted in disordered N- or C-termini, or areas with characterized ps-ns motion, labeled in blue in Fig. 4b and here. In (b,c): structures are sized by Dyna-1 p(exchange) and colored by experimental labels as in Fig. 4.

**Extended Data Figure 8:**
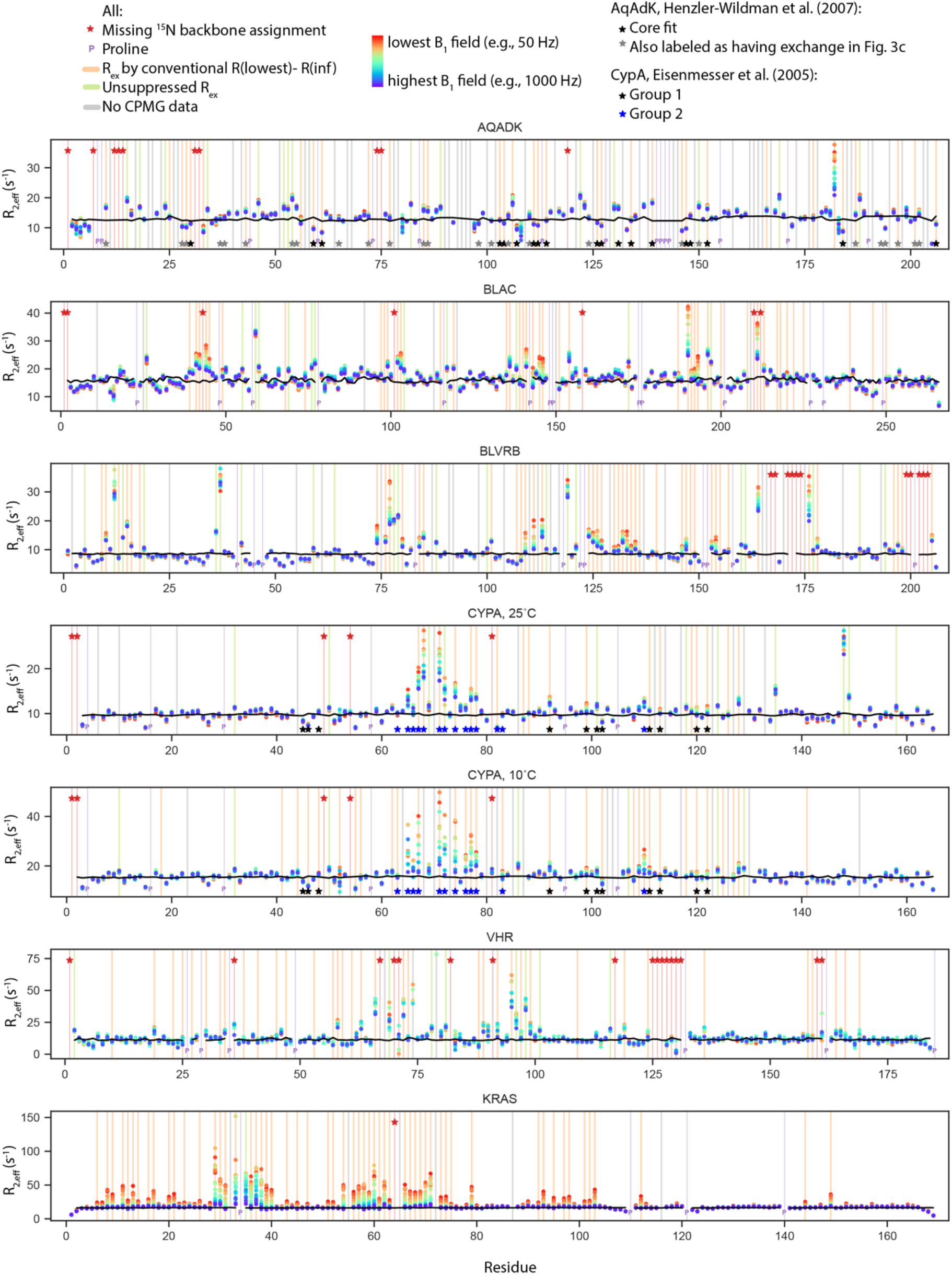
RelaxDB-CPMG datasets where dispersion data was available for all residues. Legend: Missing assignments (red star), prolines (purple “P”), exchange via R_ex_ calculated conventionally (light orange), residues with unsuppressed R_ex_ (light green), residues with no CPMG data due to peak overlap (gray). The color gradient represents B_1_ field strength from lowest (red, e.g., 50 Hz) to highest (purple, e.g., 1000 Hz). The AqAdk and CypA datasets are annotated with residue groups reported in refs. ^41^ and ^38^, respectively.

**Extended Data Figure 9:**
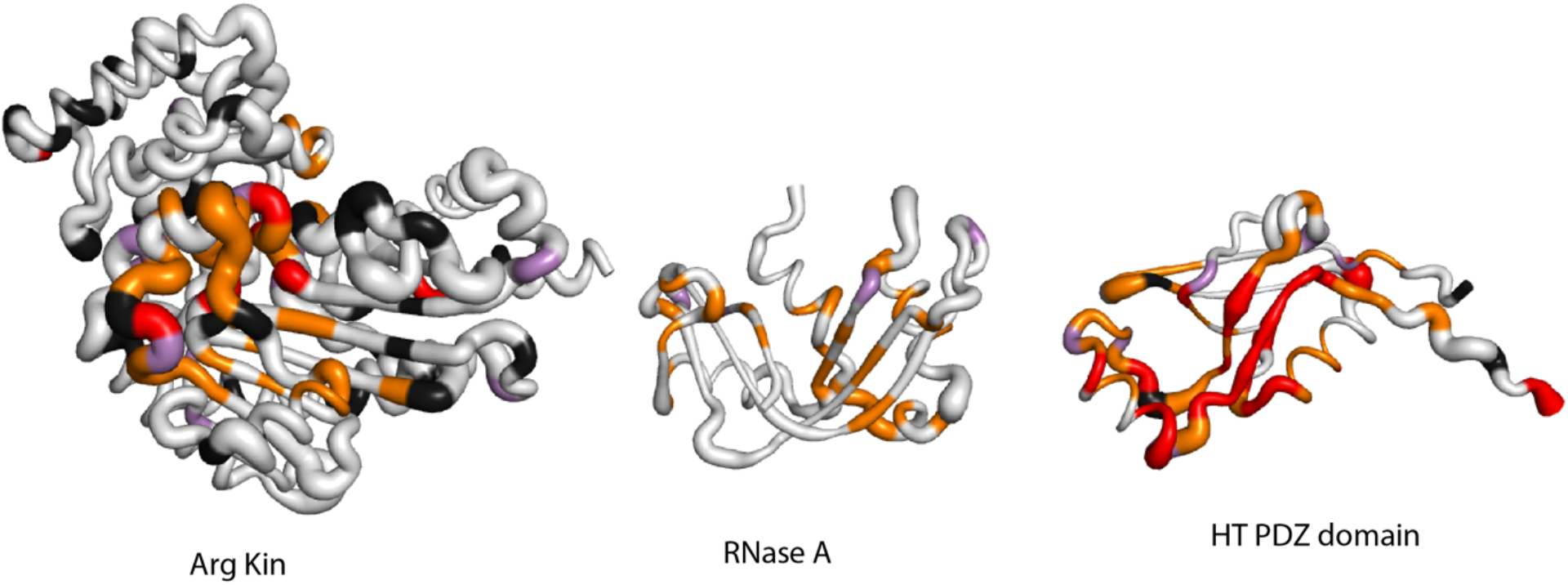
Additional ^15^N CPMG datasets with no dispersion data published for all residues (cf. Figure 5e).

## References

1 Hoch, J. C. et al. Biological Magnetic Resonance Data Bank. Nucleic Acids Res 51, D368–D376 (2023). 10.1093/nar/gkac1050

2 Hayes, T. et al. Simulating 500 million years of evolution with a language model. Science, eads0018 (2025). 10.1126/science.ads0018

3 Frauenfelder, H. Sligar, S. G. C Wolynes, P. G. The energy landscapes and motions of proteins. Science 254, 1598–1603 (1991). 10.1126/science.1749933

4 McCammon, J. A., Gelin, B. R. C Karplus, M. Dynamics of folded proteins. Nature 267, 585–590 (1977). 10.1038/267585a0

5 Jumper, J. et al. Highly accurate protein structure prediction with AlphaFold. Nature 596, 583–589 (2021). 10.1038/s41586-021-03819-2

6 Berman, H. M. et al. The Protein Data Bank. Nucleic Acids Res 28, 235–242 (2000). 10.1093/nar/28.1.235

7 Henzler-Wildman, K. C Kern, D. Dynamic personalities of proteins. Nature 450, 964–972 (2007). 10.1038/nature06522

8 van den Bedem, H. C Fraser, J. S. Integrative, dynamic structural biology at atomic resolution--it’s about time. Nat Methods 12, 307–318 (2015). 10.1038/nmeth.3324

9 Hansen, D. F. Vallurupalli, P. C Kay, L. E. Using relaxation dispersion NMR spectroscopy to determine structures of excited, invisible protein states. J Biomol NMR 41, 113–120 (2008). 10.1007/s10858-008-9251-5

10 Tollinger, M., Skrynnikov, N. R., Mulder, F. A., Forman-Kay, J. D. C Kay, L. E. Slow dynamics in folded and unfolded states of an SH3 domain. J Am Chem Soc 123, 11341–11352 (2001). 10.1021/ja011300z

11 Bruschweiler, R. New approaches to the dynamic interpretation and prediction of NMR relaxation data from proteins. Curr Opin Struct Biol 13, 175–183 (2003). 10.1016/s0959-440x(03)00036-8

12 Cilia, E., Pancsa, R., Tompa, P., Lenaerts, T. C Vranken, W. F. From protein sequence to dynamics and disorder with DynaMine. Nat Commun 4, 2741 (2013). 10.1038/ncomms3741

13 Wayment-Steele, H. K. et al. Predicting multiple conformations via sequence clustering and AlphaFold2. Nature 625, 832–839 (2024). 10.1038/s41586-023-06832-9

14 Lewis, S. et al. Scalable emulation of protein equilibrium ensembles with generative deep learning. Science 389, eadv9817 (2025). 10.1126/science.adv9817

15 Loria, J. P., Rance, M. C Palmer A. G., 3rd. A TROSY CPMG sequence for characterizing chemical exchange in large proteins. J Biomol NMR 15, 151–155 (1999). 10.1023/a:1008355631073

16 Vallurupalli, P. Bouvignies, G. C Kay, L. E. Increasing the exchange time-scale that can be probed by CPMG relaxation dispersion NMR. J Phys Chem B 115, 14891–14900 (2011). 10.1021/jp209610v

17 Clore, G. M., Driscoll, P. C., Wingfield, P. T. C Gronenborn A. M. Analysis of the backbone dynamics of interleukin-1 beta using two-dimensional inverse detected heteronuclear 15N-1H NMR spectroscopy. Biochemistry 29, 7387–7401 (1990). 10.1021/bi00484a006

18 Hall, J. B. C Fushman, D. Characterization of the overall and local dynamics of a protein with intermediate rotational anisotropy: Differentiating between conformational exchange and anisotropic diffusion in the B3 domain of protein G. J Biomol NMR 27, 261–275 (2003). 10.1023/a:1025467918856

19 Brenner, A. K., Kieffer, B., Trave, G., Froystein, N. A. C Raae A. J. Thermal stability of chicken brain alpha-spectrin repeat 17: a spectroscopic study. J Biomol NMR 53, 71–83 (2012). 10.1007/s10858-012-9620-y

20 Kim, I. et al. Solution structure and dynamics of anti-CRISPR AcrIIA4, the Cas9 inhibitor. Sci Rep 8, 3883 (2018). 10.1038/s41598-018-22177-0

21 Elings, W. et al. Two beta-Lactamase Variants with Reduced Clavulanic Acid Inhibition Display Different Millisecond Dynamics. Antimicrob Agents Chemother 65, e0262820 (2021). 10.1128/AAC.02628-20

22 Lin, Z. et al. Evolutionary-scale prediction of atomic-level protein structure with a language model. Science 379, 1123–1130 (2023). 10.1126/science.ade2574

23 Srivastava, N., Hinton, G., Krizhevsky, A. Sutskever, I. C Salakhutdinov, R. Dropout: a simple way to prevent neural networks from overfitting. The journal of machine learning research 15, 1929–1958 (2014).

24 Zhou, M. et al. in International Conference on Machine Learning. 7594–7602 (PMLR).

25 Weiss, K. Khoshgoftaar, T. M. C Wang, D. A survey of transfer learning. Journal of Big Data 3, 9 (2016). 10.1186/s40537-016-0043-6

26 Yu, V., Ronzone, E., Lord, D., Peti, W. C Page, R. MqsR is a noncanonical microbial RNase toxin that is inhibited by antitoxin MqsA via steric blockage of substrate binding. J Biol Chem 298, 102535 (2022). 10.1016/j.jbc.2022.102535

27 Basu, R., Eichhorn, C. D., Cheng, R., Peterson, R. D. C Feigon, J. Structure of S. pombe telomerase protein Pof8 C-terminal domain is an xRRM conserved among LARP7 proteins. RNA Biol 18, 1181–1192 (2021). 10.1080/15476286.2020.1836891

28 Sakhrani, V. V. et al. Backbone assignments and conformational dynamics in the S. typhimurium tryptophan synthase alpha-subunit from solution-state NMR. J Biomol NMR 74, 341–354 (2020). 10.1007/s10858-020-00320-2

29 Morin, S. C Gagne, S. M. NMR dynamics of PSE-4 beta-lactamase: an interplay of ps-ns order and mus-ms motions in the active site. Biophys J 96, 4681–4691 (2009). 10.1016/j.bpj.2009.02.068

30 Grey, M. J., Wang, C. C Palmer A. G., 3rd. Disulfide bond isomerization in basic pancreatic trypsin inhibitor: multisite chemical exchange quantified by CPMG relaxation dispersion and chemical shift modeling. J Am Chem Soc 125, 14324–14335 (2003). 10.1021/ja0367389

31 Beeser, S. A., Goldenberg, D. P. C Oas, T. G. Enhanced protein flexibility caused by a destabilizing amino acid replacement in BPTI. J Mol Biol 269, 154–164 (1997). 10.1006/jmbi.1997.1031

32 Shaw, D. E. et al. Atomic-level characterization of the structural dynamics of proteins. Science 330, 341–346 (2010). 10.1126/science.1187409

33 Baldisseri, D. M., Margolis, J. W., Weber, D. J. Koo, J. H. C Margolis, F. L. Olfactory marker protein (OMP) exhibits a beta-clam fold in solution: implications for target peptide interaction and olfactory signal transduction. J Mol Biol 319, 823–837 (2002). 10.1016/S0022-2836(02)00282-6

34 Nakashima, N., Nakashima, A. Nakashima, K. C Takano, M. Olfactory marker protein contains a leucine-rich domain in the Omega-loop important for nuclear export. Mol Brain 15, 89 (2022). 10.1186/s13041-022-00973-0

35 Salter, J. D. C Smith, H. C. Modeling the Embrace of a Mutator: APOBEC Selection of Nucleic Acid Ligands. Trends in Biochemical Sciences 43, 606–622 (2018). 10.1016/j.tibs.2018.04.013

36 Stehle, T. et al. The apo-structure of the low molecular weight protein-tyrosine phosphatase A (MptpA) from Mycobacterium tuberculosis allows for better target-specific drug development. J Biol Chem 287, 34569–34582 (2012). 10.1074/jbc.M112.399261

37 Aoto, P. C., Stanfield, R. L., Wilson, I. A., Dyson, H. J. C Wright, P. E. A dynamic switch in inactive p38γ leads to an excited state on the pathway to an active kinase. Biochemistry 58, 5160–5172 (2019).

38 Eisenmesser, E. Z. et al. Intrinsic dynamics of an enzyme underlies catalysis. Nature 438, 117–121 (2005). 10.1038/nature04105

39 Fraser, J. S. et al. Hidden alternative structures of proline isomerase essential for catalysis. Nature 462, 669–673 (2009). 10.1038/nature08615

40 Eisenmesser, E. Z., Bosco, D. A. Akke, M. C Kern, D. Enzyme dynamics during catalysis. Science 295, 1520–1523 (2002). 10.1126/science.1066176

41 Henzler-Wildman, K. A. et al. Intrinsic motions along an enzymatic reaction trajectory. Nature 450, 838–844 (2007). 10.1038/nature06410

42 Paukovich, N. et al. Biliverdin Reductase B Dynamics Are Coupled to Coenzyme Binding. J Mol Biol 430, 3234–3250 (2018). 10.1016/j.jmb.2018.06.015

43 Hansen, A. L., Xiang, X., Yuan, C. Bruschweiler-Li, L. C Bruschweiler, R. Excited-state observation of active K-Ras reveals differential structural dynamics of wild-type versus oncogenic G12D and G12C mutants. Nat Struct Mol Biol 30, 1446–1455 (2023). 10.1038/s41594-023-01070-z

44 Beaumont, V. A. et al. Allosteric Impact of the Variable Insert Loop in Vaccinia H1-Related (VHR) Phosphatase. Biochemistry 59, 1896–1908 (2020). 10.1021/acs.biochem.0c00245

45 Ohnuma, T. et al. Chitin oligosaccharide binding to a family GH19 chitinase from the moss Bryum coronatum. FEBS J 278, 3991–4001 (2011). 10.1111/j.1742-4658.2011.08301.x

46 Yee, A. et al. An NMR approach to structural proteomics. Proc Natl Acad Sci U S A 99, 1825–1830 (2002). 10.1073/pnas.042684599

47 Dudley, J. A., Park, S., MacDonald, M. E., Fetene, E. C Smith, C. A. Resolving overlapped signals with automated FitNMR analytical peak modeling. J Magn Reson 318, 106773 (2020). 10.1016/j.jmr.2020.106773

48 Garcia de la Torre, J., Huertas, M. L. C Carrasco B. HYDRONMR: prediction of NMR relaxation of globular proteins from atomic-level structures and hydrodynamic calculations. J Magn Reson 147, 138–146 (2000). 10.1006/jmre.2000.2170

49 Bernado, P., Garcia de la Torre, J. C Pons, M. Interpretation of 15N NMR relaxation data of globular proteins using hydrodynamic calculations with HYDRONMR. J Biomol NMR 23, 139–150 (2002). 10.1023/a:1016359412284

## References

1 Kneller, J. M., Lu, M. & Bracken, C. An effective method for the discrimination of motional anisotropy and chemical exchange. J Am Chem Soc 124, 1852–1853 (2002). 10.1021/ja017461k

2 Garcia de la Torre, J., Huertas, M. L. & Carrasco, B. HYDRONMR: prediction of NMR relaxation of globular proteins from atomic-level structures and hydrodynamic calculations. J Magn Reson 147, 138–146 (2000). 10.1006/jmre.2000.2170

3 Bernado, P., Garcia de la Torre, J. & Pons, M. Interpretation of 15N NMR relaxation data of globular proteins using hydrodynamic calculations with HYDRONMR. J Biomol NMR 23, 139–150 (2002). 10.1023/a:1016359412284

4 Raza, T. et al. Insights into the NF-κB-DNA Interaction through NMR Spectroscopy. ACS Omega 6, 12877–12886 (2021). 10.1021/acsomega.1c01299

5 Korn, S. M. et al. (1)H, (13)C, and (15)N backbone chemical shift assignments of the nucleic acid-binding domain of SARS-CoV-2 non-structural protein 3e. Biomol NMR Assign 14, 329–333 (2020). 10.1007/s12104-020-09971-6

6 Johnson, E. C., Lazar, G. A., Desjarlais, J. R. & Handel, T. M. Solution structure and dynamics of a designed hydrophobic core variant of ubiquitin. Structure 7, 967–976 (1999). 10.1016/S0969-2126(99)80123-3

7 Pei, J. & Grishin, N. V. AL2CO: calculation of positional conservation in a protein sequence alignment. Bioinformatics 17, 700–712 (2001). 10.1093/bioinformatics/17.8.700

8 Shrake, A. & Rupley, J. A. Environment and exposure to solvent of protein atoms. Lysozyme and insulin. J Mol Biol 79, 351–371 (1973). 10.1016/0022-2836(73)90011-9

9 McGibbon, R. T. et al. MDTraj: A Modern Open Library for the Analysis of Molecular Dynamics Trajectories. Biophys J 109, 1528–1532 (2015). 10.1016/j.bpj.2015.08.015

10 Chen, H. & Zhou, H. X. Prediction of solvent accessibility and sites of deleterious mutations from protein sequence. Nucleic Acids Res 33, 3193–3199 (2005). 10.1093/nar/gki633

11 Hoch, J. C. et al. Biological Magnetic Resonance Data Bank. Nucleic Acids Res 51, D368–D376 (2023). 10.1093/nar/gkac1050

12 Lin, Z. et al. Evolutionary-scale prediction of atomic-level protein structure with a language model. Science 379, 1123–1130 (2023). 10.1126/science.ade2574

13 Hauser, M., Steinegger, M. & Soding, J. MMseqs software suite for fast and deep clustering and searching of large protein sequence sets. Bioinformatics 32, 1323–1330 (2016). 10.1093/bioinformatics/btw006

14 Henzler-Wildman, K. A. et al. Intrinsic motions along an enzymatic reaction trajectory. Nature 450, 838–844 (2007). 10.1038/nature06410

15 Elings, W. et al. Two beta-Lactamase Variants with Reduced Clavulanic Acid Inhibition Display Different Millisecond Dynamics. Antimicrob Agents Chemother 65, e0262820 (2021). 10.1128/AAC.02628-20

16 Paukovich, N. et al. Biliverdin Reductase B Dynamics Are Coupled to Coenzyme Binding. J Mol Biol 430, 3234–3250 (2018). 10.1016/j.jmb.2018.06.015

17 Eisenmesser, E. Z., Bosco, D. A., Akke, M. & Kern, D. Enzyme dynamics during catalysis. Science 295, 1520–1523 (2002). 10.1126/science.1066176

18 Beaumont, V. A. Functional Implications of Conformational Dynamics of Protein Tyrosine Phosphatases, Yale University, (2019).

19 Hansen, A. L., Xiang, X., Yuan, C., Bruschweiler-Li, L. & Bruschweiler, R. Excited-state observation of active K-Ras reveals differential structural dynamics of wild-type versus oncogenic G12D and G12C mutants. Nat Struct Mol Biol 30, 1446–1455 (2023). 10.1038/s41594-023-01070-z

20 McDonald, L. R., Boyer, J. A. & Lee, A. L. Segmental motions, not a two-state concerted switch, underlie allostery in CheY. Structure 20, 1363–1373 (2012). 10.1016/j.str.2012.05.008

21 Aspholm, E. E., Lidman, J. & Burmann, B. M. Structural basis of substrate recognition and allosteric activation of the proapoptotic mitochondrial HtrA2 protease. Nat Commun 15, 4592 (2024). 10.1038/s41467-024-48997-5

22 Beach, H., Cole, R., Gill, M. L. & Loria, J. P. Conservation of mus-ms enzyme motions in the apo- and substrate-mimicked state. J Am Chem Soc 127, 9167–9176 (2005). 10.1021/ja0514949

23 Davulcu, O., Flynn, P. F., Chapman, M. S. & Skalicky, J. J. Intrinsic domain and loop dynamics commensurate with catalytic turnover in an induced-fit enzyme. Structure 17, 1356–1367 (2009). 10.1016/j.str.2009.08.014

24 Zhang, Y. & Skolnick, J. TM-align: a protein structure alignment algorithm based on the TM-score. Nucleic Acids Res 33, 2302–2309 (2005). 10.1093/nar/gki524

25 Case, D. A. Normal mode analysis of protein dynamics. Current Opinion in Structural Biology 4, 285–290 (1994). 10.1016/S0959-440X(94)90321-2

26 Zhang, S. et al. ProDy 2.0: increased scale and scope after 10 years of protein dynamics modelling with Python. Bioinformatics 37, 3657–3659 (2021). 10.1093/bioinformatics/btab187

27 Loshchilov, I. Decoupled weight decay regularization. arXiv preprint arXiv:1711.05101 (2017).

28 Menon, A. K. et al. Long-tail learning via logit adjustment. arXiv preprint arXiv:2007.07314 (2020).

29 Waskom, M. L. Seaborn: statistical data visualization. Journal of Open Source Software 6, 3021 (2021).

30 Yee, A. et al. An NMR approach to structural proteomics. Proc Natl Acad Sci U S A 99, 1825–1830 (2002). 10.1073/pnas.042684599

31 Taira, T. et al. Cloning and characterization of a small family 19 chitinase from moss (Bryum coronatum). Glycobiology 21, 644–654 (2011). 10.1093/glycob/cwq212

32 Hansen, D. F., Vallurupalli, P. & Kay, L. E. An improved 15N relaxation dispersion experiment for the measurement of millisecond time-scale dynamics in proteins. J Phys Chem B 112, 5898–5904 (2008). 10.1021/jp074793o

33 Jiang, B., Yu, B., Zhang, X., Liu, M. & Yang, D. A (15)N CPMG relaxation dispersion experiment more resistant to resonance offset and pulse imperfection. J Magn Reson 257, 1–7 (2015). 10.1016/j.jmr.2015.05.003

34 Ahlner, A., Carlsson, M., Jonsson, B. H. & Lundstrom, P. PINT: a software for integration of peak volumes and extraction of relaxation rates. J Biomol NMR 56, 191–202 (2013). 10.1007/s10858-013-9737-7

35 Vallurupalli, P., Bouvignies, G. & Kay, L. E. Studying “invisible” excited protein states in slow exchange with a major state conformation. J Am Chem Soc 134, 8148–8161 (2012). 10.1021/ja3001419

